# Absolute measurement of cellular activities using photochromic single-fluorophore biosensors

**DOI:** 10.1101/2020.10.29.360214

**Authors:** Vincent Gielen, Viola Mönkemöller, Franziska Bierbuesse, Anaïs C. Bourges, Wim Vandenberg, Yi Shen, Johan Hofkens, Pieter Vanden Berghe, Robert E. Campbell, Benjamien Moeyaert, Peter Dedecker

## Abstract

Genetically-encoded biosensors based on a single fluorescent protein are widely used to visualize analyte levels or enzymatic activities in cells, though usually to monitor relative changes rather than absolute values. We report photochromism-enabled absolute quantification (PEAQ) biosensing, a method that leverages the photochromic properties of biosensors to provide an absolute measure of the analyte concentration or activity. We develop proof-of-concept photochromic variants of the popular GCaMP family of Ca^2+^ biosensors, and show that these can be used to resolve dynamic changes in the absolute Ca^2+^ concentration in live cells. We also show how our method can be expanded to fast imaging with reduced illumination intensities or to situations where the absolute illumination intensities are unknown. In principle, PEAQ biosensing can be applied to other biosensors with photochromic properties, thereby expanding the possibilities for fully quantitative measurements in complex and dynamic systems.

## Introduction

Genetically encoded biosensors have revolutionized our understanding of the inner workings of the cell, by reporting on *in situ* enzymatic activities or the presence of a wide range of molecules such as ions, metabolites and messengers [1, 2]. Most of the genetically encoded biosensors can be divided into two classes, depending on whether their signal is read out as a change in Förster resonance energy transfer (FRET) between donor and acceptor fluorophores, or as a change in the fluorescence intensity of a single fluorophore [3]. These fluorophores are usually derived from fluorescent proteins (FPs), though other systems have also been used [4]. A key advantage of FRET-based probes is that they can provide absolute quantification of the analyte concentration using ratiometric readout of the FRET efficiency [5–7]. However, they suffer from complications such as incomplete FP maturation [8], take up considerable spectral bandwidth unless lifetime imaging or anistropy measurements can be used [9–11], and result in relatively large genetic constructs, which complicate genome-engineering methods such as viral gene transduction. Single FP-based biosensors, on the other hand, provide fast and straightforward recording and take up less of the visible spectrum, facilitating multiplexing with other fluorescent probes or optogenetic modulators such as channelrhodopsins [12]. They are also encoded with considerably smaller genes. However, the absolute determination of the analyte concentration or activity is usually difficult to perform with these biosensors since their emission is also affected by the local concentration of the probe and the illumination intensity, which depend on the nature of the sample and the depth of the imaging. Furthermore, these measurements provide no way to detect saturation of the biosensor response. This complicates the development of the *in-situ* quantitative biological models that are ultimately needed to understand and predict cell behavior.

Intriguingly, a variety of previous work has shown that additional information can be provided by the photophysical and photochemical properties of the fluorophore. Ratiometric excitation and/or emission biosensors, for example, provide quantitative signals where the activity is encoded as the ratio between two excitation or emission bands, and have been reported both for biosensors based on FPs [13] and organic dyes [14]. However, not all single-FP biosensors have proven amenable to ratiometric operation, while organic dyes present additional challenges for the labelling of more complex samples and when performing long-term experiments [15]. Fortunately, a range of other light-induced behaviors provide perspectives for innovative biosensor designs [16]. The development of photo-activatable and photoconvertible biosensors, such as CaMPARI [17] and CaMPARI2 [18], has enabled light-induced highlighting of specific features of a sample [17– 20]. Other strategies have built on the use of photochromic FPs, which can be reversibly switched between a fluorescent and a non-fluorescent state through irradiation with light. Their photochromism has enabled powerful technologies such as dynamic labeling and sub-diffraction imaging [21–30]. Mechanistically, this process occurs via a light-driven cis/trans isomerization of the chromophore, followed by spontaneous (de-) protonation on account of the different pK_a_’s of the cis and the trans state [31]. Within the field of biosensing, photochromism has been used to enable more accurate FRET quantification [32, 33], to switch between different FRET pairs in a single molecule or complex [34], to enable fluorescence lifetime imaging microscopy (FLIM)-like readout of the FRET activity using a conventional microscope [35], as an alternative readout of homo-FRET [36], and to multiplex spectrally-overlapping biosensors [37]. So far, however, less work has been done on the use of these properties for biosensors based on single FPs.

Ca^2+^ is a prime target for biosensing as it is a crucial secondary messenger in many cellular processes such as neuronal signaling, muscle contraction and the regulation of cell fate. The desire to record its spatiotemporally dynamic levels in situ [38, 39] has led to the development of genetically-encoded Ca^2+^ indicators (GECIs) based on single FPs, which translate the intracellular Ca^2+^ concentration into a fluorescent signal [40]. Using these tools, Ca^2+^ concentrations can be monitored in a variety of systems, ranging from single cells to the brain [41]. However, these studies are often focused on relative changes in local Ca^2+^ levels, as the quantitative assessment of these signals is limited by the intensiometric nature of single-FP GECIs, though a number of ratiometric variants have also been developed, including Ratiometric-Pericam [42], GEM-GECO and GEX-GECO [43]. Alternative ratiometric systems have also been pursued by fusing a second, red-shifted fluorescent protein to an intensiometric GECI, but this results in a construct that shares the limitations associated with FRET-based GECIs [15]. A range of organic dyes, such as the Fura and Indo families [14, 44] also achieve ratiometric readout.

Even though single FP-based biosensors are often treated as exhibiting fluorescence that is solely dependent on the free analyte concentration, previous investigations reported that the emissive signal can also depend on the previous light irradiation [17, 45]. Recent work also showed that it is possible to engineer Ca^2+^ sensitivity into an existing photochromic protein to obtain an erasable integrator of neuronal activity [46]. We reasoned that photochromism might form a more general mechanism to obtain additional information from genetically encoded biosensors.

In this work, we set out to develop a strategy for determining absolute analyte activities using photochromic single FP-based biosensors. First, we developed proof-of-concept GECIs that display photochromism that is highly dependent on the local Ca^2+^ concentration. We then demonstrate that this dependence can be used to determine the absolute Ca^2+^ concentration in cells in a way that is largely insensitive to photobleaching and the intensity of excitation light. By delivering fully quantitative information using single FP-based biosensors, we expect that our approach can be used to perform quantitative biosensing across a variety of systems.

## Results and Discussion

### Design of photochromic GECIs

The GCaMP family is the most widely used class of single FP-based Ca^2+^ biosensors [47]. They consist of a circularly permutated variant of EGFP and a Ca^2+^-sensitive moiety made up of a calmodulin (CaM) domain and RS20, a CaM-binding peptide. Binding of Ca^2+^ ions by CaM leads to an interaction between CaM and RS20, resulting in a fluorescence increase that is largely due to a change in the pK_a_ of the chromophore [48–50]. Well-performing photochromic variants of EGFP were previously reported [26, 51–53], which suggested that photochromic behavior might be introduced in the GCaMP biosensor design while retaining its sensitivity to Ca^2+^.

We selected GCaMP3 [54], GCaMP6 (-s, -m, and -f variants) [55], GCaMP7s [47], and GECO1.1 and GECO-1.2 [43] as the starting scaffolds, and introduced mutations that were shown to induce photochromism in EGFP, in particular T65A, Q69L, V150A, V/A163S, and S205N (numbering according to EGFP) [26, 51]. The resulting proteins were expressed in *E. coli* and screened for brightness and reversible photochromism with cyan and violet light (Supplementary Figures S1 to S3). We found that the single point mutation T65A or Q69L sufficed to introduce pronounced photochromism into the GECIs, while still retaining the sensitivity to Ca^2+^, though at the cost of a reduction in overall fluorescence brightness in bacterial lysates. We attempted to improve this brightness by introducing several mutations (S30R, Y39N, N105Y, and I171V) that were previously shown to increase the expression levels of EGFP [52, 56] and GCaMP variants [57, 58]. However, these so-called superfolder mutations did not appear to influence the overall brightness in bacterial lysates and had a negative impact on the photochromism and Ca^2+^ sensitivity (Supplementary Figure S1). We therefore excluded these variants from further consideration. Based on their brightness and photochromism, we identified GCaMP3-T65A (GCaMP3-T) and GCaMP3-Q69L (GCaMP3-Q), and GCaMP6s-T and -Q as the most promising mutants, together making up a new family of photochromic genetically-encoded Ca^2+^ indicators.

We next characterized their spectroscopic properties *in vitro* (Figure 1, Table 1 and Supplementary Figures S4 to S9). Overall, the absorption and fluorescence spectra are similar, although the GCaMP3-T and GCaMP6s-T variants show a ∼10 nm blue-shift in their absorption, excitation, and emission maxima. We expressed the Ca^2+^-sensitivity of the fluorescence using (ΔF/F_*sat*_)_Ca_, the difference in the observed fluorescence between the Ca^2+^-saturated (*sat*) and Ca^2+^-free states (*apo*), normalized by the fluorescence of the *sat*-form, resulting in Ca^2+^ sensitivities that are in line with those reported in literature [55] (Figure 1C and Supplementary Figure S1).

**Figure 1:**
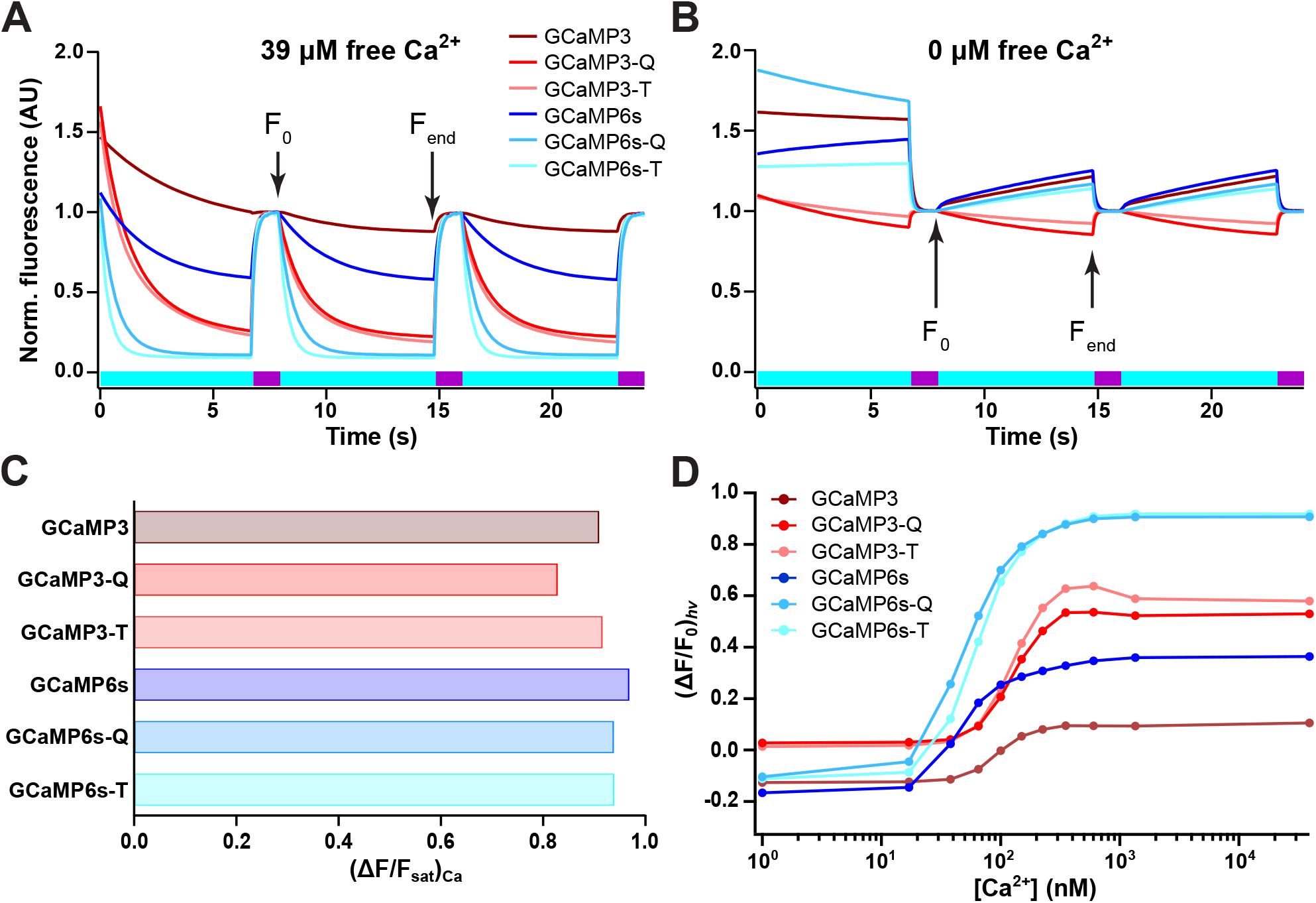
*In vitro* dependence of the sensor fluorescence on light illumination and Ca^2+^ levels. (**A**-**B**) Cyan- and violet-light-induced photochromism in purified proteins in the presence (**A**) and absence (**B**) of free Ca^2+^. The cyan and violet bars at the bottom of the plots show the sequence of the illumination. F_0_ and F_end_ refer to the intensities that would be used to calculate the (ΔF/F_0_)_hν_ values. (**C**) Difference in the observed fluorescence between Ca^2+^-saturated (*sat*) and Ca^2+^-free conditions (*apo*), normalized by the fluorescence of the *sat*-form, as measured in bacterial cell lysates. (**D**) Photochromism contrast (ΔF/F_0_)_hν_ as a function of [Ca^2+^].

**Table 1:** Spectroscopic properties of GCaMP3, GCaMP6s and their -Q and -T mutants. In the case where two values are present (e.g. 403/503), the first and second values are associated with the neutral and anionic forms of the chromophore, respectively. The extinction coefficients are ‘apparent’ in that they are expressed with respect to the total protein concentration, rather than purely the concentration of the fluorescent state, and are reported for the absorption maxima. Spectra, pH titrations and Ca^2+^ titrations are available in Supplementary Figures S1 to S9.

|  | GCaMP3 | GCaMP3-Q | GCaMP3-T | GCaMP6s | GCaMP6s-Q | GCaMP6s-T |
| --- | --- | --- | --- | --- | --- | --- |
| $\lambda_{\text{abs}, \text{apo}}$ (nm) | 403/503 | 404/503 | 397/495 | 400/502 | 400/510 | 395/493 |
| $\lambda_{\text{abs}, \text{sat}}$ (nm) | 396/499 | 400/498 | 394/486 | 401/497 | 405/496 | 399/486 |
| $\lambda_{\text{exc}, \text{apo}}$ (nm) | 397/504 | 401/505 | 387/491 | 398/511 | 393/512 | 392/495 |
| $\lambda_{\text{exc}, \text{sat}}$ (nm) | 498 | 497 | 391/484 | 497 | 496 | 483 |
| $\lambda_{\text{em}, \text{apo}}$ (nm) | 515 | 515 | 510 | 518 | 518 | 512 |
| $\lambda_{\text{em}, \text{sat}}$ (nm) | 513 | 513 | 506 | 513 | 513 | 506 |
| $\epsilon_{\text{apo}}^{\text{app}}$ ($10^3 \text{ M}^{-1} \text{ cm}^{-1}$) | 2.4 | 6.7 | 3.8 | 8.0 | 5.7 | 5.2 |
| $\epsilon_{\text{sat}}^{\text{app}}$ ($10^3 \text{ M}^{-1} \text{ cm}^{-1}$) | 19.0 | 24.3 | 13.4 | 63.7 | 67.5 | 48.0 |
| QY of fluorescence | 0.59 | 0.29 | 0.10 | 0.60 | 0.34 | 0.07 |
| Molecular brightness (vs EGFP) | 0.33 | 0.21 | 0.035 | 1.13 | 0.69 | 0.083 |
| $\text{pK}_{\text{a}, \text{sat}}$ | 7.6 | 7.6 | 8.1 | 5.9 | 6.0 | 6.2 |
| $\text{pK}_{\text{a}, \text{apo}}$ | 10.10 | 8.4 | 8.7 | 12.7 | 10.2 | 11.8 |
| $K_{\text{d}}$ (nM) ($\text{Ca}^{2+}$, fluorescence) | 191 | 177 | 238 | 181 | 266 | 261 |
| $K_{\text{d}}$ (nM) ($\text{Ca}^{2+}$, $(\Delta F/F_0)_{\text{hv}}$) | 95 | 123 | 118 | 47 | 51 | 63 |
| Hill coefficient | 3.1 | 3.1 | 3.3 | 2.1 | 2.1 | 2.3 |
| $(\Delta F/F_{\text{sat}})_{\text{Ca}}$ (on) | 0.90 | 0.71 | 0.89 | 0.98 | 0.97 | 0.92 |
| $(\Delta F/F_{\text{sat}})_{\text{Ca}}$ (off) | 0.89 | 0.47 | 0.55 | 0.97 | 0.74 | -0.11 |
| $(\Delta F/F_0)_{\text{hv}}$ (*sat*) | -0.19 | 0.65 | 0.83 | 0.082 | 0.86 | 0.93 |
| $(\Delta F/F_0)_{\text{hv}}$ (*apo*) | -0.44 | 0.39 | 0.31 | -0.32 | 0.0071 | -0.010 |
| Thermal recovery, *sat* (s) | 110 | 308 | 2388 | 38 | 330 | 726 |

The photochromism of each probe was then assayed in 11 different free Ca^2+^ concentrations ranging from 0 to 39 μM. The GCaMP3-T, GCaMP3-Q, GCaMP6s-T and GCaMP6s-Q mutants clearly show enhanced photochromism over their parent proteins (Figure 1A,B and Supplementary Figure S8), where the photochromism contrast (F_0_ − F_end_)/F_0_ = (ΔF/F_0_)_hν_ strongly depends on the free Ca^2+^ concentration (Figure 1D). We obtained different *K*_*d*_ values for the fluorescence and the (ΔF/F_0_)_hν_ titration (Table 1), though both titrations could be well described using a conventional sigmoidal fit. While a full analysis is outside of the scope of this contribution, this difference reflects that the photochromism involves the contributions of and interconversions between multiple states, whereas the F_0_ titration probes only the presence of the fluorescent (deprotonated cis) state. A more detailed investigation of the dynamics of photochromic sensors was recently published in Ref. [59].

Interestingly, the (ΔF/F_0_)_hν_ values are negative at low, but positive at high Ca^2+^ concentrations for GCaMP3, GCaMP6s, GCaMP6s-Q and GCaMP6s-T (Figure 1D and Supplementary Figure S8). This reflects that these proteins show positive photochromism at low Ca^2+^ concentrations (cyan excitation light leads to an *increase* in fluorescence), and negative photochromism at higher Ca^2+^ concentrations (cyan excitation light leads to a *decrease* in fluorescence). This change in photochromism modalities likely results from changes in the pK_a_’s of the fluorescent (cis) and non-fluorescent (trans) chromophore states due to Ca^2+^ binding, since these pK_a_’s determine the directionality of the photochromism [60].

The photochromism data (Figure 1A,B) also reveal that the first photochromic cycle differs from all subsequent cycles, even though the illumination patterns are identical. This phenomenon is present to a variable degree in all photochromic fluorescent proteins [61, 62]. Successful photochromism-based sensing using such labels thus requires that an initial photochromism cycle is always performed at the beginning of an experiment, and discarded from the analysis. A few more cycles may need to be discarded if the photochromism cycles are incomplete due to lower-intensity or short illumination, which can be determined by inspecting when the photochromism reaches a stable level. Left in the dark, the samples recover back to the initial state following the spontaneous thermal recovery pathway, which usually takes on the order of minutes or longer (Table 1).

Overall, our *in vitro* characterization revealed that the GCaMP6s variants had an approximately 3-fold higher molecular brightness (Table 1) and an up to 9-fold higher Ca^2+^ response (F_*sat*_ − F_*apo*_, Supplementary Figure S1) in comparison to the GCaMP3 variants. They also show a slightly higher photochromic contrast and a reduced pH sensitivity. Moreover, the GCaMP3-Q and GCaMP6s-Q variants have a 8- to 10-fold higher molecular brightness compared to the GCaMP3-T and GCaMP6s-T variants. As a result of this overall analysis, we selected GCaMP6s-Q as the best variant for our further experiments. We also established that the photochromic behaviour of this probe shows very little temperature dependence between room temperature and 37°C (Supplementary Figure S10), and also tested the fatigue resistance of GCaMP6s-Q at 39 μM Ca^2+^ both in solution and in permeabilized cells, finding that it retains about 50% of the original fluorescence after 1000 photochromism cycles, while the (ΔF/F_0_)_hν_ value is much less affected by this repeated photochromism. A similar measurement in calcium-free conditions likewise showed that the (ΔF/F_0_)_hν_ metric is largely robust against changes in the probe fluorescence (Supplementary Figure S11). Furthermore, these *in vitro* and in-cell measurements revealed highly similar kinetics for the photochromism in vitro and in cells (Supplementary Figure S12).

In summary, our measurements indicate that our photochromic sensors can be represented using the scheme shown in Figure 2, consisting of four different states characterized by whether Ca^2+^ is bound and whether the molecule is in the spectroscopic on- or off-state.

**Figure 2:**
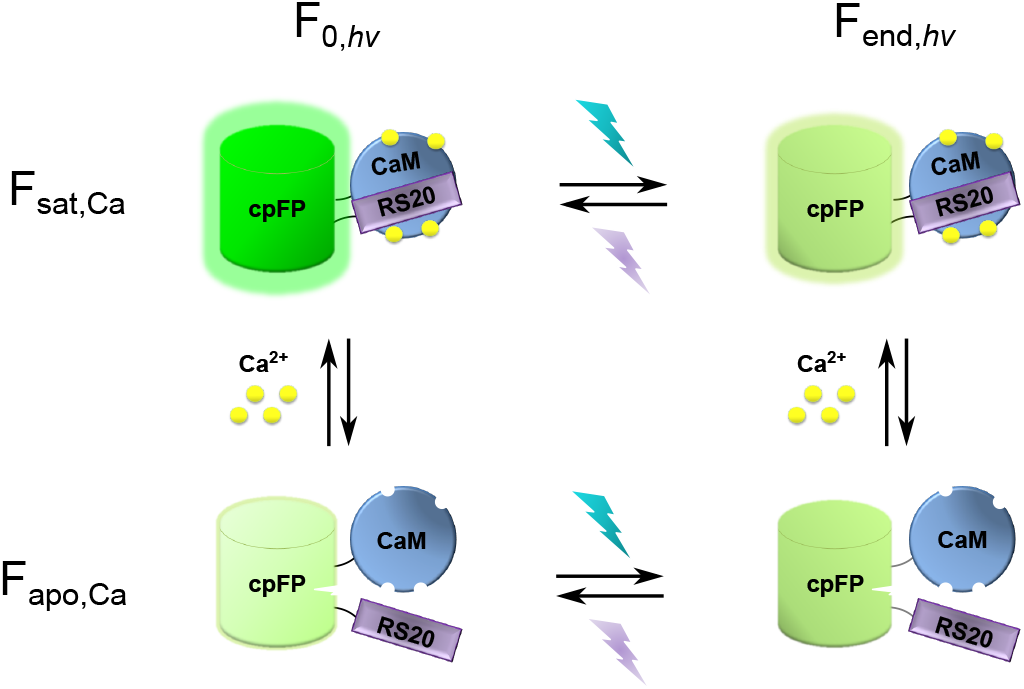
Schematic model for a single-FP-based photochromic sensor, involving the presence of four different states as determined by the absence or presence of Ca^2+^ and the spectroscopic on- or off-state. In the *sat* state the cyan and violet light cause off- and on-switching of the fluorescence, respectively, which is reversed in the *apo* state.

### PEAQ biosensing estimates absolute Ca^2+^ concentrations in live cells

We next expressed GCaMP6s-Q in HeLa cells and measured its photochromism at different Ca^2+^ concentrations by immersing the cells in Ca^2+^-buffered solution supplemented with ionomycin and saponin. The individual cells show pronounced photochromic responses (Supplementary Figure S13A), though displaying significant cell-to-cell variability. We found that most or all of this variability was due to the low brightness of GCaMP6s-Q-expressing cells, suggesting a reduced expression, maturation, and/or increased degradation in HeLa cells (Supplementary Figure S13B–C).

These results suggested that our method could be used to dynamically measure absolute Ca^2+^ concentrations inside the brighter live cells. To verify this, we stimulated GCaMP6s-Q-transfected Hela cells with 25 μM histamine and measured a photochromism cycle every 2 seconds (Figure 3A), obtaining absolute fluorescence intensity and derived (ΔF/F_0_)_hν_ traces for each cell. Using the *in vitro* calibration curve measured using the same instrumental settings (Figure 3C), we could determine the absolute Ca^2+^ concentration at each point in the measurement (Figure 3B-D). Though the histamine-induced oscillations display high cell-to-cell variability, the magnitude of these concentrations is similar to those observed in previous work [63, 64]. We further compared the Ca^2+^ concentrations obtained upon histamine stimulation using this approach and the FRET-based indicator Yellow Cameleon 2.60 [65], resulting in very similar values (Supplementary Figure S14).

**Figure 3:**
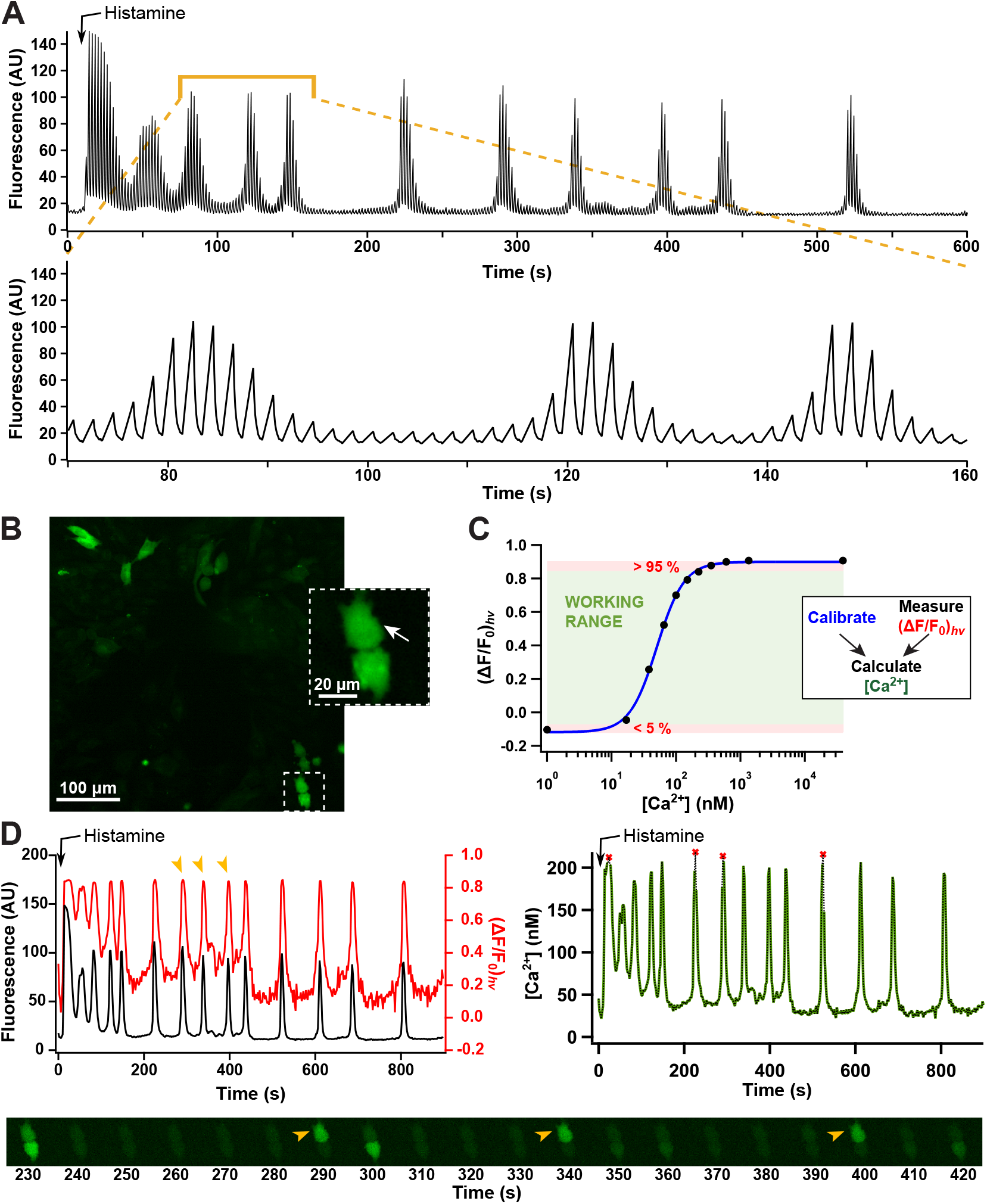
(**A**) Fluorescence time trace of a GCaMP6s-Q-expressing HeLa cell after stimulation with 25 μM histamine, recorded by performing a photochromism cycle every two seconds. The off-switching light also serves as the excitation light for imaging. (**B**) Fluorescence images of representative HeLa cells expressing GCaMP6s-Q. The expansion and white arrow mark the cell for which the PEAQ analysis is shown in panel D. (**C**) (ΔF/F_0_)_hν_ calibration curve at different [Ca^2+^], measured *in vitro* using the same instrumental settings as for cell imaging. The extreme 5% regions (red shading) are considered to be out of the working range (green shading). The inset to panel C shows a schematic overview of the methodology. (**D**) Fluorescence (black), (ΔF/F_0_)_hν_ (red), and absolute [Ca^2+^] (green, right panel) traces of the cell indicated with the white arrow in panel B after stimulation with 25 μM histamine. Red markers represent [Ca^2+^] calculated from (ΔF/F_0_)_hν_ values in the extreme 5% range of the calibration curve. The image sequence shows the change in fluorescence throughout a subset of the experiment. Specific time points are highlighted in the graphs and images (orange arrows).

We decided to name our method photochromism-enabled absolute quantification (PEAQ) biosensing. In principle, the PEAQ methodology can be combined with any biosensor where the presence of the sensed stimulus results in a change in photochromic properties. The low brightness of cells expressing GCaMP6s-Q restricted our analysis to regions-of-interest consisting of an entire cell, though the methodology could be applied to smaller regions or even in a pixelwise manner when combined with sufficiently bright signals. Since it is based on the quantification of fluorescence intensities, our method is sensitive to the presence of background signal, though this sensitivity is highly similar to that observed for FRET-based quantification (Supplementary Note 1).

We opted to use the *in vitro* calibration as it is obtained in well-defined conditions for which the Ca^2+^ concentration is unambiguously known. The presence of cell-to-cell variability has also led to the development of *in situ* calibration schemes, in which the biosensors are driven to the unbound and fully-bound state post-measurement by replacement of the medium with Ca^2+^-free and high [Ca^2+^] buffers containing ionophores and/or permeabilizing agents [5, 49, 66]. The resulting dynamic range of the signal is then used to correct the per-cell response. Our methodology should be readily compatible with this approach since the response of our system displays a similar saturation behavior in sufficiently bright live cells (Supplementary Figure S13). However, in either case the calibration should be done using instrument settings matching those used in the actual experiment.

Our results allow us to not only estimate the absolute Ca^2+^ concentration, but also to infer when saturation of the biosensor occurs, a distinction that would be more difficult to make using purely intensiometric readout. We used this to visually indicate (ΔF/F_0_)_hν_ values that fell into the upper 5% signal range of the *in vitro* calibration curve (Figure 3C), where this cutoff was chosen based on visual inspection of the curve. In principle, a similar consideration applies also to the lower 5% signal range, though low Ca^2+^ concentrations are more difficult to measure accurately due to the lower fluorescence emission. These 5% thresholds indicate conditions in which large change in Ca^2+^ results in only a small change in (ΔF/F_0_)_hν_, rendering such measurements less reliable for direct interpretation.

A single PEAQ acquisition requires the measurement of a fluorescence image, irradiation with off-switching light, the measurement of a second fluorescence image, and irradiation with on-switching light. For this reason, the temporal resolution of the method is considerably lower compared to more standard intensiometric measurements that require just a single fluorescence measurement. The required irradiation duration is determined by the speed of the switching, which in turn depends on the intensity of the irradiation and the switching propensity of the probe. Using our instrument, we determined the time constants of the photochromism to be approximately 0.18 s for the calcium-saturated and 0.50 s for the calcium-free form. With our setup and the described measurement protocol, we can achieve a temporal resolution of up to 1.1 s for one PEAQ acquisition. However, we do note that it is possible to mitigate this reduced temporal resolution using an approach based on combined photochromism and intensiometric measurements, as we will discuss in the next section of this work.

The temporal resolution of PEAQ can also be adapted by choosing the duration of the off-switching irradiation after which F_end_ is measured. Shorter durations can lead to considerably faster measurements, but may also affect the measurement uncertainty since the photochromism is less pronounced. We chose to set the irradiation such that the fluorescence approached the plateau for the calcium-saturated state, though a very similar performance can be achieved using considerably less irradiation (Supplementary Figure S15). In any case, a change in F_end_ also requires a corresponding change in the construction of the calibration curve (Supplementary Figure S16).

Because our readout is ratiometric and does not depend on the absolute fluorescence intensities, it is less susceptible to photobleaching or dynamic changes in biosensor concentration, as was already shown *in vitro* (Supplementary Figure S11). Figure 3D, for example, shows gradually decreasing fluorescence levels that likely reflect the onset of photodestruction. However, the insensitivity of our photochromism-based readout to fluorophore concentration is reflected in the essentially constant amplitude of the (ΔF/F_0_)_hν_ oscillations.

### Combined photochromic and intensiometric measurements enable lower light doses and faster acquisitions

A possible downside of PEAQ biosensing is that a full photochromism cycle must be acquired for each absolute measurement (Figure 4A). Switching the fluorophores to the off-state requires more time and more excitation light compared to performing a regular fluorescence acquisition, and could result in a reduced temporal resolution and more photodestruction or phototoxicity.

**Figure 4:**
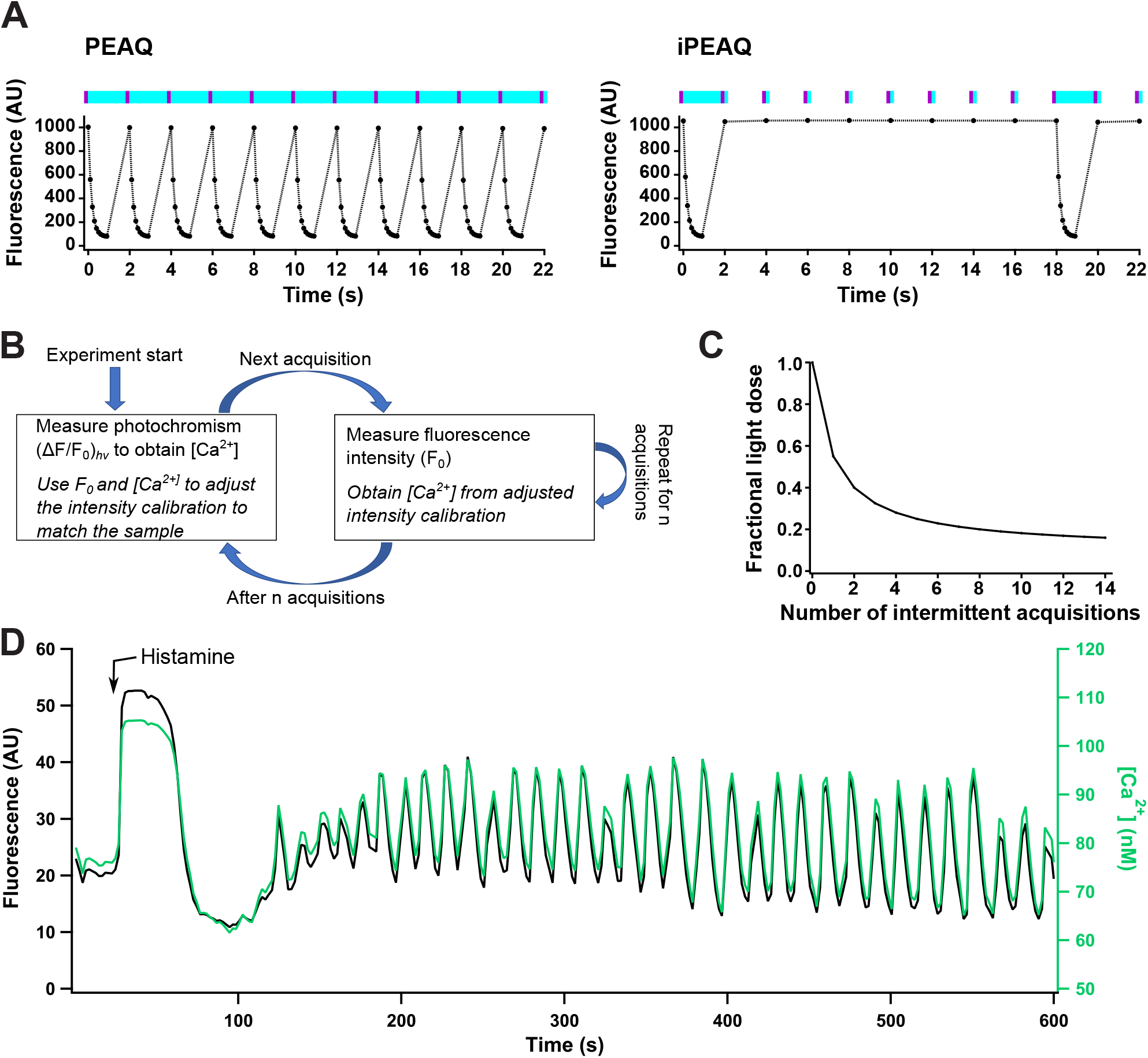
iPEAQ biosensing with intermittent calibration. (**A**) Comparison of the irradiation procedure of PEAQ biosensing using repeated photochromism cycles (left), and iPEAQ biosensing using intermittent photochromism cycles (right). (**B**) Schematic overview of the intermittent calibration methodology. (**C**) Fraction of the light dose that can be saved using iPEAQ as a function of the number of intensiometric acquisitions per photochromism cycle. (**D**) Fluorescence intensity F_0_ (black) and [Ca^2+^] trace (green) of a single GCaMP6s-Q-expressing HeLa cell after stimulation with 25 μM histamine (black).

A key aspect of our probes is that the Ca^2+^ concentration affects both the kinetics of the photochromism as well as the overall fluorescence brightness of the probe (Supplementary Figures S8 and S9). We reasoned that the cyan light dose could be reduced by alternating the on/off photochromism cycles with standard acquisitions of single fluorescence images. Ordinarily, such fluorescence images do not allow the estimation of the absolute analyte concentration since the relation between the fluorescence and the Ca^2+^ concentration depends on the probe concentration. However, we have shown above that the (ΔF/F_0_)_hν_ signal can be translated into the absolute Ca^2+^ concentration (Supplementary Figure S17). Knowledge of the Ca^2+^ concentration and corresponding fluorescence brightness can then be used to adjust the fluorescence-Ca^2+^ titration to match the specifics of the sample, allowing the determination of absolute Ca^2+^ values based only on fluorescence measurements. This suggests a light-efficient and fast approach to measure the absolute Ca^2+^ concentration: an initial photochromism cycle is performed, yielding an absolute Ca^2+^ concentration via the (ΔF/F_0_)_hν_ calibration. This Ca^2+^ concentration is then used to scale the intensity calibration curve to match the brightness of the sample. Subsequent measurement points then measure only the fluorescence brightness. Assuming that the local probe concentration does not change, any changes in the observed fluorescence must be due to changes in the Ca^2+^ concentration and can be translated into an absolute Ca^2+^ concentration using the known [Ca^2+^]-intensity relationship. A more precise description of the methodology is given in the materials and methods section of this work.

A conceptual overview of this method is shown in Figure 4B. The resulting impact on the cyan light dose is pronounced, with the total cyan light dose reducing to less than half if (ΔF/F_0_)_hν_ acquisitions are alternated with just two regular fluorescence acquisitions (Figure 4C). The violet light dose before each acquisition can also be correspondingly reduced since less off-switching also means that less light is required to recover the biosensors to the fluorescent state. Finally, because measuring a single fluorescence image is faster than measuring a full off-switching event, this approach could also be used to quantitatively measure transient dynamics that occur too quickly to permit a full photochromism-based acquisition.

In principle, the photochromism-based calibration could be performed just once at the beginning of the acquisition, with single fluorescence acquisitions used for every timepoint thereafter. In practice, however, the local probe concentration may vary due to the dynamics of the sample, leading to fluorescence intensity changes that are not due to changes in analyte activity. A more robust approach is therefore to perform regular photochromism-based acquisitions alternated with a fixed number of fluorescence acquisitions, which we describe as ‘intermittent PEAQ’ (iPEAQ), where the duration between photochromism-based acquisitions should be sufficiently short that changes in the local probe concentration are small or negligible. However, photochromic sensors that show different dissociation constants for photochromism and intensiometric measurements may limit the analyte concentration window over which the calibration can be performed successfully.

We verified the feasibility of this approach on HeLa cells expressing GCaMP6s-Q stimulated with histamine. Figure 4D and Supplementary Figure S18 show the results for a repeated sequence of eight fluorescence acquisitions between every photochromism-based calibration. Supplementary Figure S19 shows how very similar peak Ca^2+^ levels can be extracted from cells showing very different cell brightnesses. This procedure allowed us to reduce the cyan light exposure by 80% while still delivering direct measurements of the absolute Ca^2+^ concentration.

### PEAQ biosensing does not require knowledge of the absolute illumination intensities

The extent of the sensor photochromism depends not only on the Ca^2+^ concentration, but also on the intensity and duration of the cyan light. In many systems the exact excitation intensity at the sample may not be known precisely, such as when performing measurements deeper in tissues, where part of the light may be absorbed or scattered before reaching the region of interest. Such conditions also preclude the quantitative determination of the activity using purely intensiometric measurements, even if the probe concentration were known. We therefore investigated whether our method could be used reliably when the absolute excitation intensity is unknown.

Our key observation is that the fluorescence cannot be suppressed completely through irradiation with off-switching light, but instead reaches a plateau that reflects an equilibrium between the on- and off-switching of the fluorophore. Because both the initial fluorescence intensity and the plateau scale with the applied light intensity (Figure 5A), the resulting (ΔF/F_0_)_hν_ values are largely independent of the excitation intensity (Supplementary Figure S20). Provided that the applied light dose is sufficient to approach the plateau, which can be verified by monitoring the evolution of the fluorescence in time, the resulting measurements do not require knowledge of the absolute illumination intensities. Alternatively, the level of the plateau can be extrapolated by fitting each of the acquired photochromism cycles with an exponential function (Figure 5B). To verify this approach, we measured photochromism in 11 different Ca^2+^ concentrations in a representative HeLa cell at 47 mW and 24 mW excitation power (Figure 5C). Values for (ΔF/F_0_)_hν_ were independent of the method of choice and yielded very similar Ca^2+^ concentrations over a range of excitation intensities (Figure 5C and Supplementary Figure S20). This finding makes it possible to apply our method also in more complex samples.

**Figure 5:**
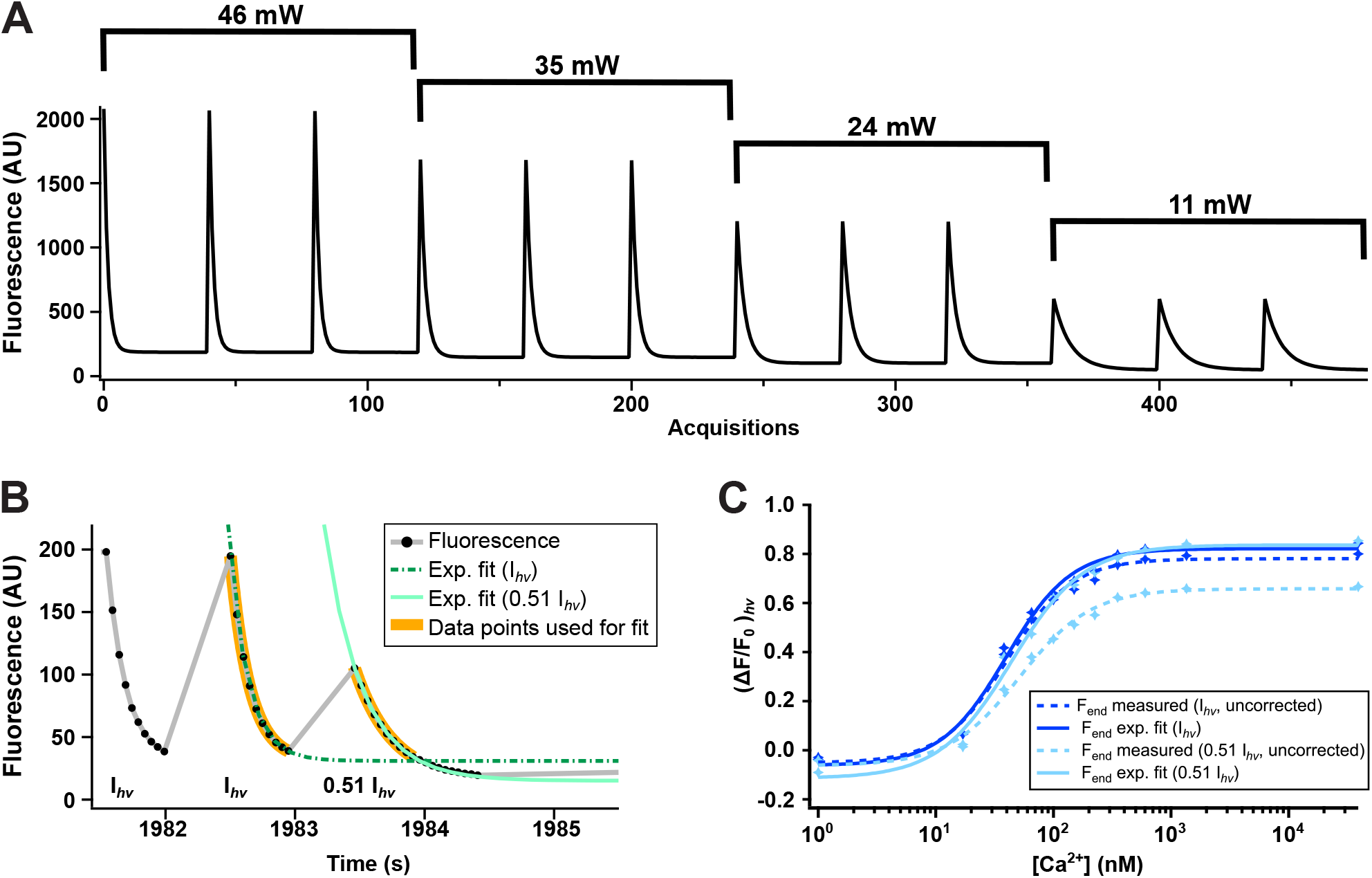
Intensity-independence of PEAQ biosensing. **A**: Raw fluorescence traces obtained on purified GCaMP6s-Q protein at 39 μM free Ca^2+^ with varying powers of cyan irradiation. Full data including other Ca^2+^ concentrations is shown in Supplementary Figure S20. **B**: Raw fluorescence trace (black) of a representative GCaMP6s-Q-expressing Hela cell at high Ca^2+^ concentration during photoswitching. Off-switching was achieved via either 10 acquisitions with 47 mW cyan excitation (I_h*v*_), or 20 acquisitions with 24 mW cyan excitation (0.51 I_h*v*_). Green dashed lines show the exponential fits of the off-switching decays. The thick orange bands indicate the 10 data points used for fitting. **C**: Ca^2+^ titration of GCaMP6s-Q expressed in a representative HeLa cell. (ΔF/F_0_)_hν_ was calculated using either F_end_ measured after ten acquisitions, or an F_end_ parameter obtained from the plateau of an exponential fit. ‘Uncorrected’ indicates calibration curves that were obtained without exponential fitting. For other curves the F_end_ signal was determined using the exponential fit-based procedure.

## Conclusions

We have demonstrated a novel approach for the imaging of absolute cellular activities using photochromic biosensors, which we call photochromism-enabled absolute quantification (PEAQ) biosensing. Starting from the GCaMP family of Ca^2+^ biosensors, we showed how pronounced photochromism could be introduced using a limited set of mutations. We found that the single point mutations T65A or Q69L were sufficient to gain a 10-fold improvement in photochromism contrast, and determined that these mutants retained their strong sensitivity to the local Ca^2+^ concentration. We found that the combination of GCaMP6s with Q69L yielded the most promising performance *in vitro*, and named this mutant GCaMP6s-Q.

The Ca^2+^ concentration not only affected the overall fluorescence brightness but also altered the photochromic properties of the biosensor. We quantified the extent of the photochromism by measuring the fluorescence before and after illumination with cyan light, providing a metric that is conceptually straightforward and easily measured. *In vitro* calibration of (ΔF/F_0_)_hν_ versus [Ca^2+^] allowed us to determine the absolute concentration of Ca^2+^ in solution.

We then demonstrated the measurement of intracellular Ca^2+^ concentrations in live HeLa cells stimulated with histamine, enabling us to follow a diverse range of Ca^2+^ dynamics. Our approach not only allowed the measurement of absolute Ca^2+^ concentrations, but also made it possible to determine when these results are less reliable due to saturation of the biosensor, and is independent of changes in local probe concentration that may occur through sample dynamics or photodestruction. We did find that GCaMP6s-Q displayed poor brightness when expressed in cells, indicating that further probe development will be needed to achieve robust performance across a variety of systems.

A possible downside of PEAQ biosensing is that inducing the photochromism involves more irradiation and is slower compared to the acquisition of a single fluorescence image, increasing the chance of phototoxic effects and making it more difficult to resolve fast dynamics. In response, we developed iPEAQ, a scheme in which quantitative photochromism-based measurements are alternated with one or more regular fluorescence acquisitions. Ordinarily, such intensity-based acquisitions do not permit the quantitative determination of the biosensor activity. However, we showed that absolute quantification can be achieved for both types of acquisitions, by leveraging the dependence of both the photochromism and the fluorescence brightness on the Ca^2+^ concentration. In principle, a single photochromism-based reference measurement is sufficient to enable fast quantitative measurements for the entire duration of the experiment, though in practice periodic photochromism-based measurements are likely required to adapt to the dynamics of the sample. The ability to introduce a non-invasive reference measurement at any time is a key advantage of our method. An iPEAQlike procedure could also readily be used in other situations where an orthogonal readout is available, as would be the case when measuring the analyte using e.g. fluorescence lifetime imaging.

Many experimental scenarios require measurements in more complex samples such as tissues, where the absolute intensity of the excitation light is typically not known due to the absorption and scattering. We find that our method does not require knowledge of the absolute excitation intensity, provided that the measurements are performed in such a way that the photochromism is allowed to go to a level where the on- and off-switching are in equilibrium, or to a level where this equilibrium can be estimated. This can be readily verified, even if the absolute intensity is unknown, by monitoring the fluorescence response to the illumination. This approach is also fully compatible with the reduced light doses afforded by our alternating photochromism- and intensity-based measurements, and should allow our method to scale to a broad variety of samples and settings.

As it utilizes a single FP-based biosensor and ratiometric readout, our method is also less sensitive to issues such as limited FP maturation or photodestruction, while also being readily amenable to the multiplexing of multiple biosensors emitting in different spectral bands. We also expect that our methodology can be applied to any FP-based biosensor that displays, or can be engineered to display, photochromism, including the many GCaMP-like biosensors that have been developed. Overall, our work readily extends the possibilities for fully quantitative measurements inside complex systems.

## Acknowledgements

We thank Sam Duwé, Wim Vandenberg, Hideaki Mizuno (KU Leuven) and Alison Tebo (Janelia Research Campus) for critical comments and insightful discussion. We also thank Hideaki Mizuno for providing us with the Yellow Cameleon YC2.60 plasmid. We thank Beatrice Adelizzi, Thomas le Saux, and Ludovic Jullien (ENS Paris) for critical comments and insightful discussion related to the acquisition methodology and data analysis. We thank Thijs Roebroek (KU Leuven) for assistance with the imaging. We thank Abhi Aggarwal (University of Alberta) for technical assistance and helpful discussion. V.G. thanks the Research Foundation Flanders (FWO Vlaanderen) for a doctoral fellowship. A.B. thanks the European Commission for a Marie Skłodowska-Curie fellowship. B.M. thanks the Research Foundation Flanders for a postdoctoral fellowship. Work in the lab of R.E.C. was supported with funding from the University of Alberta and the Natural Sciences and Engineering Research Council of Canada (RGPIN-2018-04364). This work was supported through funding from the Research Foundation Flanders through grants 1514319N, G090819N, G0B8817N, and the European Research Council through grant 714688 NanoCellActivity.

## Author contributions

P.D. designed and supervised research. V.G., V.M., B.M., A.B., F.B., and P.D. designed experiments. V.G., V.M., A.B., F.B., and B.M. performed experiments. V.M., F.B., and W.V. performed analysis of the cell-based experiments. V.G., V.M., F.B., B.M., and W.V. performed analysis of the*in vitro* experiments. Y.S. identified the T65A mutation and edited the manuscript. R.E.C. supervised research on the T65A mutation and edited the manuscript. P.V.B. contributed ideas, discussions and advice concerning ratiometric Ca^2+^ measurements in live cells. J.H. contributed critical discussion. P.D., V.G., V.M., B.M., and F.B. wrote the manuscript with input from all authors.

## Methods

### Cloning and Site-Directed Mutagenesis

For expression in *E. coli*, GCaMP3, 6s, 6m, 6f, 7s, GECO1.1 and GECO1.2 were cloned in a pRSET B vector and mutations were introduced using a modified QuikChange protocol [67]. Eukaryotic expression vectors were created by inserting PCR-amplified GECI genes between BamHI and EcoRI restriction sites of a pcDNA3 vector. GCaMP6s-Q was also cloned from pCDNA3 into the chicken beta-actin promotor vector (which appears to have increased protein expression) [9]. Primers were ordered at Integrated DNA Technologies and all constructs were verified by DNA sequencing (LGC Genomics).

### Bacterial Cell Lysate Screening

The GECIs were transformed into JM109(DE3) competent cells (Promega) and incubated at 37°C. The next morning, colonies were inoculated into 1.2 mL of LB media supplemented with ampicillin in 96 deep-well blocks. Cultures were grown at 30°C for 36 h, then harvested by centrifugation at 4000 rpm for 20 minutes at 4°C, frozen, thawed, resuspended in 500 μL lysis buffer (100 mM MOPS, 100 mM KCl, 1 mg/mL lysozyme, pH 7.4), and subsequently incubated at 30°C for 1 h while shaking. Cell debris was pelleted by centrifugation at 4000 rpm, and 95 μL clarified lysate was transferred to a 96-well fluorescence microplate (Greiner). The cell lysate samples were mixed with EGTA to a final concentration of 1 mM, and fluorescence was measured at 490 nm excitation and 530 nm emission (Tecan Safire II, bandwith 20 nm, gain 40 V). Subsequently, CaCl_2_ (5 mM final concentration) was added to the lysates and the fluorescence was measured again.

The photochromism was measured on an Olympus IX71 inverted microscope equipped with a Spectra X Light Engine (Lumencor), a 10× UplanSApo objective (Olympus), a ZT488RDC dichroic mirror and ET525/30m emission filter (both Chroma) and an ORCA-Flash4.0 LT+ sCMOS camera (Hamamatsu). Lysates were switched off in 30 steps of 250 ms with cyan light (470/24 nm, 32 mW, corresponding to 453 mW/cm^2^) and switched back on in 30 steps of 50 ms with violet light (395/25 nm, 12 mW, corresponding to 170 mW/cm^2^). After each step an image was acquired with a camera exposure time of 20 ms and 10 mW cyan excitation light (470/24 nm, 141 mW/cm^2^).

The sensors with the T65A mutation were discovered during an independent and parallel effort. This effort aimed to produce a sensor by grafting the photochromism-inducing mutations of rsEGFP and rsEGFP2 [26, 51] onto the GFP-based GECI, GCaMP6s [55]. Towards this end, a bacterial expression plasmid pBAD-GCaMP6s was first created by PCR amplification of the GCaMP6s gene from pGP-CMV-GCaMP6s (Addgene Plasmid #40753), followed by a Gibson Assembly (New England Biolabs) reaction with the PCR product and an XhoI/HindIII-digested pBAD/His-B vector (Thermo Fisher Scientific). Using pBAD-GCaMP6s as the template, five separate primers, each encoding a mutation from rsEGFP or rsEGFP2 (T65A, Q69L, V150A, V163S, or S205N, numbered according to EGFP), were pooled together and used to perform multisite mutagenesis using the QuikChange Multi site-directed mutagenesis kit (Agilent). Due to imperfect reaction efficiency, this procedure was expected to introduce all five single mutations and all possible combinations of multiple mutations. The resulting plasmid library was used to transform DH10B electro-competent cells (Thermo Fisher Scientific). Transformed bacteria were plated on LB-agar plates and incubated overnight at 37°C. A total of 48 green fluorescent colonies were picked from the plates and cultured in 5 mL LB media with ampicillin and arabinose at 37°C overnight. Bacteria were pelleted by centrifugation and soluble cell lysates were extracted using Bacterial Protein Extraction Reagent (B-PER, Thermo Fisher Scientific). Lysates were then screened for photochromism. Briefly, bacterial lysate samples in clear PCR tubes were illuminated for 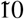 seconds using a custom-built bacterial colony fluorescence imaging macroscope [68] to either switch-on (violet light, 405/40 nm) or switch-off (cyan light, 470/40 nm) the protein fluorescence. The fluorescence change was assessed by visual inspection, and samples exhibiting photochromic behavior were immediately transferred to a multi-well plate to record the fluorescence excitation and emission spectrum using a fluorescence plate reader (Tecan). Of the 48 colonies screened, two photochromic variants were identified: GCaMP6s-T and GCaMP6s-Q. Later effort focused on the GCaMP6s-Q mutant due to its more pronounced photochromism contrast.

### Spectroscopy of Purified Proteins

Proteins were purified largely following our previously described method [52]. The purified proteins were diluted in 30 mM MOPS, 100 mM KCl, pH 7.2 (Calcium Calibration Buffer #1, C3008MP, Thermo Fisher Scientific) containing either 10 mM CaEGTA (39 μM Free Calcium Buffer) or 10 mM EGTA (Zero Free Calcium Buffer). Fluorescence excitation and emission spectra were measured on an Edinburgh 920FLS fluorimeter with 1.5 nm slits. Excitation spectra were taken from 300 to 515 nm, with emission at 520 nm. Emission spectra were taken from 485 to 700 nm, with excitation at 480 nm. Spectra were normalized to the calcium-saturated excitation and emission maxima, respectively. Absorption spectra were taken on a UV/VIS spectrometer (Ocean Optics USB4000, Ocean Insight), and normalized using the absorbance at 280 nm.

### *In vitro* Ca^2+^ Titrations

Ca^2+^ affinity assays were performed by employing a reciprocal dilution method with the Calcium Calibration Buffer Kit #1, according to the manufacturer’s instructions (C3008MP, Thermo Fischer Scientific). A small aliquot of the stock protein solution was added to 2 mL of the Zero Free Ca^2+^ Buffer (10 mM EGTA, 100 mM KCl, 30 mM MOPS, pH 7.2) to create the “zero Ca^2+^ sample.” The “high Ca^2+^ sample” was prepared by diluting exactly three times as much protein into 6 mL of 39 μM Free Ca^2+^ Buffer (10 mM CaEGTA, 100 mM KCl, 30 mM MOPS, pH 7.2). The initial 0 mM CaEGTA/GCaMP sample was loaded in the cuvette, and absorbance and emission spectra were recorded. Consecutively, the measured sample was used to prepare the next solution by removing a given volume from the sample and replacing this with an equal aliquot of the “high Ca^2+^ sample”.

For the *in vitro* Ca^2+^ titration, the same small amount of the stock protein was added to 1.2 ml of the Zero Free Ca^2+^ Buffer and to 1.2 mL of the 39 μM Ca^2+^ Buffer. The two buffer solutions were mixed in a 96 well plate to a final volume of 200 μL in order to obtain the desired EGTA concentrations from 10 mM to 0 mM. The plate was stored at 4 ° C and measured the next morning on the Olympus IX71 microscope (ZT488RDC, ET525/30m, 10× UplanSApo, Lumencor Spectra X light source) at room temperature to evaluate the Ca^2+^ dependence of (ΔF/F_0_)^hν^. We subjected each well to 15 off/on photochromic cycles. From these, the (ΔF/F_0_)^hν^ was calculated as (F_0_ - F_end_)/F_0_ with F_0_ the fluorescence read at the first frame of the second off-switching cycle and F_end_ the last frame of the second off-switching cycle (Figure 1A and B). The proteins were submitted to 15 photochromism cycles in which they were switched on in one irradiation step of 100 ms with violet light (395/25 nm, 59.3 mW, corresponding to 839 mW/cm^2^) and switched off in 10 image acquisition steps of 100 ms exposure time with cyan light (470/24 nm, 46.7 mW, 661 mW/cm^2^).

The *in vitro* titrations for the analysis of the PEAQ and iPEAQ measurements were measured with the same irradiation procedure and parameters as used during the actual measurements. The amount of cycles was reduced to 15 cycles per Ca^2+^ concentration.

### Spectroscopic characterization

Spectroscopic properties of the sensor variants were investigated in buffer solutions in the absence (*apo*) or presence (*sat*) of free Ca^2+^. Further spectral analysis (absorbance and emission) of Ca^2+^ titrations, and respectively, photochromism, thermal recovery, and the quantum yield of fluorescence, all in *apo* and *sat*, were measured on a home-built setup described elsewhere [69].

The GCaMP6s-Q photochromism *in vitro* fatigue was measured on the Olympus IX71 microscope (ZT488-RDC, ET525/30m, 10× UplanSApo, Lumencor Spectra X light source). Purified GCaMP6s-Q protein was diluted in 39 μM Free Calcium Buffer and Zero Free Calcium Buffer and transferred to a 96-well plate. The well plate was covered to prevent evaporation. The protein was switched off with ten acquisitions (100 ms cyan irradiation, 470/24 nm, 46.7 mW, about 661 mW/cm^2^), and switched back on with one 100 ms violet irradiation (395/25 nm, 59.3 mW, 839 mW/cm^2^). This was repeated every 2 s for 1000 cycles to determine the decrease in on-state fluorescence (F_0_) and photochromism contrast (ΔF/F_0_)_hν_ after repeated photochromism.

For the *in vitro* kinetics, the well plate was prepared as described before for the fatigue measurements and measured on the Olympus IX71 microscope (ZT488RDC, ET525/30m, 10× UplanSApo, Lumencor Spectra X light source). The protein was completely switched off in 30 steps of 200 ms cyan irradiation (470/24 nm, 16.1 mW, about 227 mW/cm^2^), and switched back on in 30 steps of 10 ms violet irradiation (395/25 nm, 13.7 mW, 194 mW/cm^2^). This cycle was repeated 3 times. Images were recorded after each irradiation with 9.6 mW cyan excitation (470/24 nm, 136 mW/cm^2^) and 100 ms exposure time.

Extinction coefficients (*ϵ*_*apo*_ and *ϵ*_*sat*_) were determined according to Ward’s method[70] using the literature values of EGFP, GCaMP3 and -6s as references. Quantum yield (QY) was determined relative to EGFP, having a QY of 0.6 [71]. Molecular brightness was defined as the product of *ϵ*_*sat*_ and QY, scaled to a value of 100 for EGFP.

pH titrations were performed in a universal buffer consisting of 50 mM citrate, 50 mM Tris, 50 mM glycine, 100 mM NaCl, and either 5 mM CaCl_2_ (pK_a, *sat*_) or 5 mM EGTA (pK_a, *apo*_). Absorbance spectra were measured on a Varioskan LUX Multimode Microplate Reader at pH values ranging from 4.5 to 10.5 and a plot of the absorption maxima of the anionic form in function of the pH was fitted to a sigmoid curve to yield the pK_a_.

### Cell Culture

HeLa cells cells were cultured in DMEM supplemented with 10% FBS, glutaMAX, and 0.1% gentamicin (all Gibco) at 37°C and 5% CO_2_. Before transfection, cells were seeded in 35 mm glass bottom dishes (MatTek), and allowed to settle between 4 h and 24 h. Cells were transfected using the FuGENE 6 Transfection Reagent (Promega) at a ratio of 1 μg DNA/3 μL FuGENE 6 according to the manufacturer’s protocol. Cells were supplied with fresh growth medium after 24 h, and imaged 48 h after transfection. The estimated transfection efficiency was around 80%.

For the co-culture of GCamp6sQ and Yellow Cameleon YC2.6, Hela cells were independently transfected in two flasks (25 ml) using the aforementioned transfection protocol. After 30 h, the flasks were rinsed twice with PBS and then the cells were detached using 500 μl of PBS with 0.05% trypsin. After 5 minutes of incubation at 37 °C, 4.5 ml of DMEM supplemented with FBS and glutaMAX was added. Cells transfected with GCamp6sQ and Yellow Cameleon YC2.6 were mixed in a 3:1 ratio. Then 600 μl of the resulting cell mixture was added to 35 mm dishes containing 1 ml of DMEM supplemented with FBS and glutaMAX.

### Microscopy in Mammalian Cells

For imaging, cells were rinsed twice and maintained in Hanks’ Balanced Salt Solution (HBSS; Invitrogen), supplied with 20 mM HEPES and 2 g/L D-glucose at pH 7.4 (HHBSS). Comparable solutions were made without Ca^2+^ and Mg^2+^, but with all other components (HHBSS(-)). All imaging experiments were performed on the Olympus IX71 microscope (ZT488RDC, ET525/30m, 10× UplanSApo, Lumencor Spectra X light source) at room temperature previously described.

For the Ca^2+^ titration in HeLa cells, the cells were washed twice and kept in HHBSS(-). Subsequently, a 2× stock solution of EGTA/ionomycin was added to the cells to a final concentration of 3 mM EGTA and 5 μM ionomycin, followed by incubation at room temperature for 10 minutes. Then, the buffer was replaced with ionomycin- (5 μM) and saponin- (0.005%) supplemented Zero Free Ca^2+^ Buffer, and acquisition was started immediately. Three photochromism cycles were recorded every 90 seconds. The medium’s Ca^2+^ concentration was changed every 180 seconds through a reciprocal dilution of 10 mM CaEGTA buffer (supplemented with 5 μM ionomycin and 0.005% saponin) into the Zero Free Ca^2+^ Buffer, according to the manufacturer’s protocol (Ca^2+^ Calibration Buffer Kit #1). The proteins were switched on using a single 50 ms pulse of violet light (395/25 nm, 59.3 mW, corresponding to 839 mW/cm^2^), while off-switching was achieved by acquiring 10 images with 46.7 mW of cyan excitation light (470/24 nm, 661 mW/cm^2^) and a camera exposure time of 100 ms. For the cell titrations, multiple 35mm glass-bottom dish samples were imaged at two sample positions each.

For the in cell fatigue measurement, cells were rinsed twice with HHBSS without Ca^2+^ and incubated for 10 min at room temperature with Zero Free Ca^2+^ buffer supplemented with 5 μM of ionomycin. The buffer was replaced with the same solution with 0.005% saponin for the acquisition of the *apo* state fatigue measurement or the 39 μM Ca2+ buffer supplemented with 5 μM ionomycin and 0.005% saponin for the *sat* state. The in cell fatigue measurement was conducted with the aforementioned *in vitro* imaging parameters.

The cell kinetics measurements were performed with the same imaging parameters as described in the *in vitro* spectroscopic characterization. The cells were permeabilized using the Zero Free or 39 μM Free Ca^2+^ buffers supplemented with 10 μM rotenone, 5 μM cyclopiazonic acid, 1.8 μM 2-deoxy-D-glucose and 10 μM 4-bromo-A23187 according to the protocal described in [72]. The measurements were repeated multiple times with the same dish at different sample positions.

For the imaging of histamine-induced Ca^2+^ dynamics, cells were imaged for a duration of 15 minutes during which the GCaMP6s-Q was switched on (100 ms pulse of 59.3 mW violet light, corresponding to 839 mW/cm^2^, 395/25 nm) and off (10 acquisitions, 46.7 mW cyan light, corresponding to 661 mW/cm^2^, 470/24 nm, 100 ms exposure time) every 2 s. Cells were washed twice with regular HBSS, and approximately 15 s after the start of the experiment, histamine (50 μL) was added to the cells to a final concentration of 25 μM. For each independent histamine stimulation experiment, a 35mm glass-bottom dish sample was imaged at one sample position. Each experiment was repeated with multiple dishes over multiple days. To determine [Ca^2+^] during histamine-induced dynamics, a (ΔF/F_0_)_hν_ titration curve has to be measured *in vitro* under exactly the same settings as for the actual experiments.

The intermittent PEAQ (iPEAQ) measurements were conducted analogously to the PEAQ experiments as described in the previous paragraph. PEAQ acquisitions were alternated with eight fluorescence acquisitions. The PEAQ experimental parameters can be found in the previous paragraph. The eight fluorescence acquisitions were performed with a 100 ms pulse of 59.3 mW violet light (corresponding to 839 mW/cm^2^, 395/25 nm) to switch GCaMP6s-Q on, followed by one image acquisition with a 100 ms cyan light pulse (46.7 mW, corresponding to 661 mW/cm^2^, 470/24 nm). These intensiometric acquisitions were repeated every 2 s.

For the histamine-induced co-culture experiments with GCaMP6s-Q (cloned into chicken beta-actin promotor vector) and Yellow Cameleon YC2.60, the described PEAQ measurement protocol for histamine-induced Ca^2+^ dynamics was extended by a FRET measurement. Two acquisitions with 100 ms blue light (35.6 mW, 504 mW/cm^2^, 440/20 nm, Chroma T455LP dichroic mirror) for the donor (Chroma ET480/40m emission filter) and acceptor (Chroma ET545/40m emission filter) were acquired, as well as a detection with teal light (100 ms, 7.52 mW, 106 mW/cm^2^, 510/25 nm, Chroma ZT514RDC dichroic mirror) to register the acceptor bleaching (Chroma ET545/40m emission filter). The combined PEAQ and FRET measurement scheme was repeated every 10 s. To determine [Ca^2+^], a (ΔF/F_0_)_hν_ titration curve and FRET titration curve has been measured *in vitro* under exactly the same settings as for the actual experiments, for both GCaMP6s-Q and Yellow Cameleon YC2.60. Only cells that showed a clear response and a peak brightness above 30 camera counts (in both donor and acceptor channels for Cameleon) were included in the analysis. Cameleon-expressing cells that did not show a clear anticorrelation between the donor and acceptor emission channels were excluded.

### Power independence

Measurements *in vitro* and in HeLa cells were performed as described previously. Cyan powers used for excitation were 10.9 mW, 23.8 mW, 35.4 mW, and 46.2 mW as measured at the top of the objective. The measured powers correspond to power densities of 154 mW/cm^2^, 337 mW/cm^2^, 501 mW/cm^2^ and 654 mW/cm^2^.

### Data analysis

All data analysis was done using IgorPro 8.0 (Wavemetrics). Titration curves were fitted with the following Hill equation:

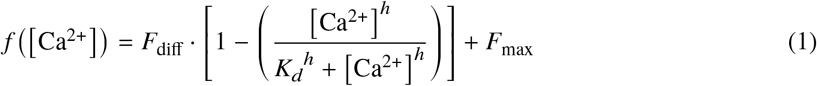

where *f* ([Ca^2+^]) is the measured (ΔF/F_0_)_hν_ or fluorescence intensity, *F*_max_ is the maximal value, *F*_diff_ is the difference between the the minimal and maximal value, *h* is the Hill coefficient, and *K*_*d*_ is the dissociation constant.

A background correction was applied to HeLa cell data, by subtracting the average intensity trace of a sample area containing no cells.

#### Analysis of the Ca^2+^ titration in HeLa cells

F_0_ and F_end_ were obtained from background-corrected average intensity traces of selected cells, and were used to calculate the (ΔF/F_0_)_hν_ values. Titration curves of the individual cells are plotted as (ΔF/F_0_)_hν_ as a function of [Ca^2+^].

#### Analysis of the histamine-induced Ca^2+^-oscillations

F_0_ and F_end_ were obtained from background-corrected average intensity traces of selected cells, and were used to calculate (ΔF/F_0_)_hν_ values. In the graphs shown in Figure 3D, the black curve (fluorescence intensity) corresponds to F_0_, and the red curve corresponds to (ΔF/F_0_)_hν_. The [Ca^2+^] (green curve) is calculated by using the *in vitro* titration curve measured using the exact same settings. Values in the lowest and highest 5% of the titration curve are defined as out of the working range (as explained in Figure 3), and are therefore plotted as red dots. Values within the working range are shown in green.

#### iPEAQ imaging

The analysis of iPEAQ data requires *in vitro* Ca^2+^ titrations acquired with the aforementioned iPEAQ measurement protocol. Two separate titrations can be extracted for the PEAQ measurements ((ΔF/F_0_)hν) and the intermittent fluorescence acquisitions (see Figure 4A). We fitted the Hill equation (1) to both titrations, resulting in two sets of parameters, one for the PEAQ and one for the fluorescence data. The fitted parameters for the fluorescence titration were then normalized so that the brightness values ranged from zero to one. We denote the resulting normalized fluorescence titration curve as *F*_*T*_ ([Ca2+]). Let us also denote the inverse calibration curve, that provides the Ca2+ concentration corresponding to a particular normalized fluorescence intensity, as 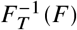.

In a first step, all of the photochromism measurements (calibration points) are extracted from the background-corrected cell fluorescence traces. The corresponding [Ca^2+^] at each measurement is then determined from the observed (ΔF/F_0_)_hν_ and the corresponding PEAQ titration parameters. Because we know the fluorescence brightness F_0_ and Ca^2+^ concentration at each calibration point, we simply determine a scale factor *A* such that

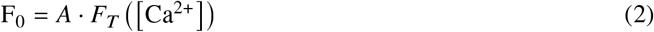

From this analysis we therefore obtain a set of {*A*_*i*_} factors, one for each calibration point *i*, that are proportional to the local probe concentration.

At the intermittent measurement points we record only F_0_ rather than a full photochromism cycle. At each such point we can then recover the local Ca^2+^ concentration using

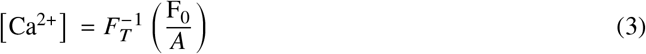

which provides the Ca^2+^ concentrations at the intermittent points where no photochromism is measured. In principle, the *A* values should be constant at each calibration measurement point if the probe concentration does not change in time. In practice, we found that these showed deviations in time due to the low brightness of our probe in cells (Supplementary Figure S21). Accordingly, we fitted these values to a line, where the standard deviation (uncertainty) of each point was set to the reciprocal of the observed fluorescence brightness (1/F_0_), and interpolated the *A* value used to determine the local Ca^2+^ concentration in eqn. 3 from this linear fit. We furthermore found that the first fluorescence acquisition after each switching cycle showed slight deviations, most likely due to the formation of a transient non-fluorescent intermediate that is not fully relaxed within 2 s, but does disappear after 4 s. This effect could be corrected through reference measurements if required, but we have instead opted to replace this first point with a linearly interpolated value between the preceding and following data points since this did not appreciably change the resulting Ca^2+^ traces.

### Data availability

The datasets generated during and/or analysed during the current study are available from the corresponding author on reasonable request.

### Code availability

The code used during the current study is available from the corresponding author on reasonable request.

