## Supplementary information for "Absolute measurement of cellular activities using photochromic single-fluorophore biosensors"

#### Supplementary note 1: effect of fluorescence background

Our approach is sensitive to background fluorescence because it relies on the quantification of absolute fluorescence signals. In the presence of background $b$ our quantification becomes

$$(\Delta\mathrm{F}/\mathrm{F}_0)_{\mathrm{h}\nu} = \frac{(F_0 + b) - (F_{\mathrm{end}} + b)}{F_0 + b} \tag{1}$$

$$= \frac{F_0 - F_{\mathrm{end}}}{F_0 + b} \tag{2}$$

from which we see that the background will affect the magnitude of the $(\Delta\mathrm{F}/\mathrm{F}_0)_{\mathrm{h}\nu}$signal. It also follows that, when doing background correction, underestimating the background (not subtracting all of it) will lead to an artificially reduced signal, while the opposite will lead to an artificially increased signal.

How sensitive is the $(\Delta\mathrm{F}/\mathrm{F}_0)_{\mathrm{h}\nu}$ to changes in the background? To answer that question we can take the derivative:

$$\frac{\mathrm{d}\,(\Delta\mathrm{F}/\mathrm{F}_0)_{\mathrm{h}\nu}}{\mathrm{d}b} = \frac{F_{\mathrm{end}} - F_0}{(b + F_0)^2} \tag{3}$$

which shows (as expected) that the distortion due to background reduces as the probe becomes brighter.

We can also ask how this compares with the signal of a ratiometric dye, such as a FRET-based $\mathrm{Ca}^{2+}$ sensor. The signal of such a system is often quantified using the emission ratio $R$:

$$R = \frac{F_A + b}{F_D + b} \tag{4}$$

where $F_A$ and $F_D$ are the signals of the acceptor and donor, and we assume for simplicity that both channels contain a background contribution $b$. To evaluate the sensitivity of the signal to background, we can likewise take the derivative:

$$\frac{\mathrm{d}R}{\mathrm{d}b} = \frac{F_D - F_A}{(b + F_D)^2} \tag{5}$$

which shows that background sensitivity of our method is in fact the same as that found for the ratiometric imaging of FRET sensors.

We also examined how the background affects the $(\Delta F/F_0)_{h\nu}$ values obtained in actual measurements. Supplementary Figure S22A shows the $(\Delta F/F_0)_{h\nu}$ values calculated for the data in main text Figure 3D and E, where each trace corresponds to a different $(\Delta F/F_0)_{h\nu}$ measurement at various $Ca^{2+}$ levels. The figure shows the effect of adding or subtracting counts to the raw data from the camera, as is done during a correction for background. This shows that the $(\Delta F/F_0)_{h\nu}$ becomes smaller when adding background counts and larger when subtracting them, as is also consistent with equation (2).

The data analysis used in this work assumes that the background $b$ can be estimated as the signal coming from a cell-free region, about 130 camera counts using our instrument. Supplementary Figure S22B shows an expansion of this region in panel A. Measurements on untransfected cells show that this autofluorescence is around 4 counts on average, rendering it likely that our estimated $b$ is slightly too low. As this expansion shows, this does result in a small change in $(\Delta F/F_0)_{h\nu}$. In principle, we could also subtract this additional background as well, though we opted for the simplicity of the cell-free subtraction.

### Supplementary Figures

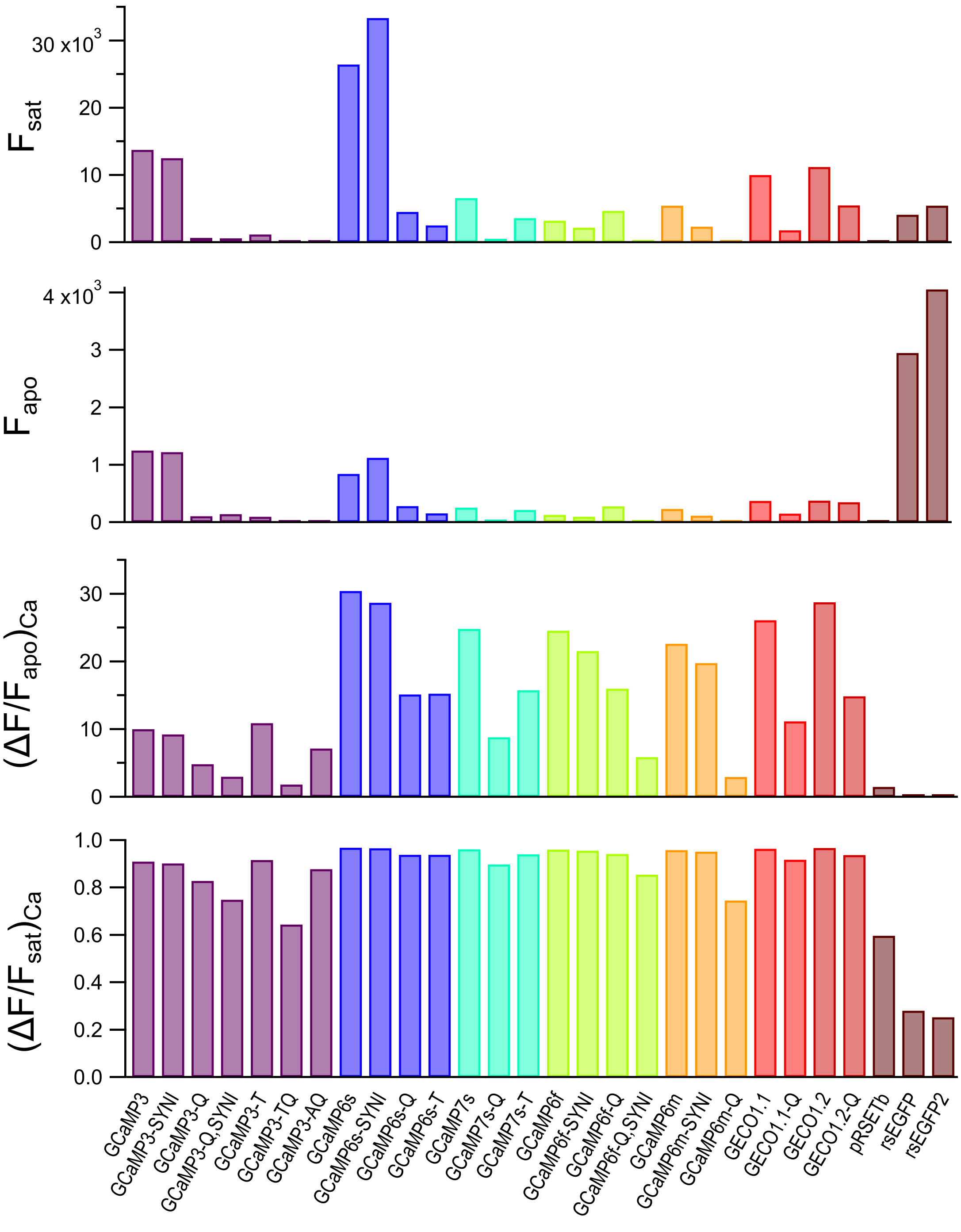


Supplementary Figure 1: Characterization of all mutants in bacterial cell lysates. Values are the averages of $n = 2$ or $n = 3$ lysates. (A) Fluorescence in the presence of 5 mM $CaCl_2$. (B) Fluorescence in the presence of 1 mM EGTA ($Ca^{2+}$-free). (C) $(\Delta F/F_{apo})_{\text{Ca}}$. (D) $(\Delta F/F_{sat})_{\text{Ca}}$.

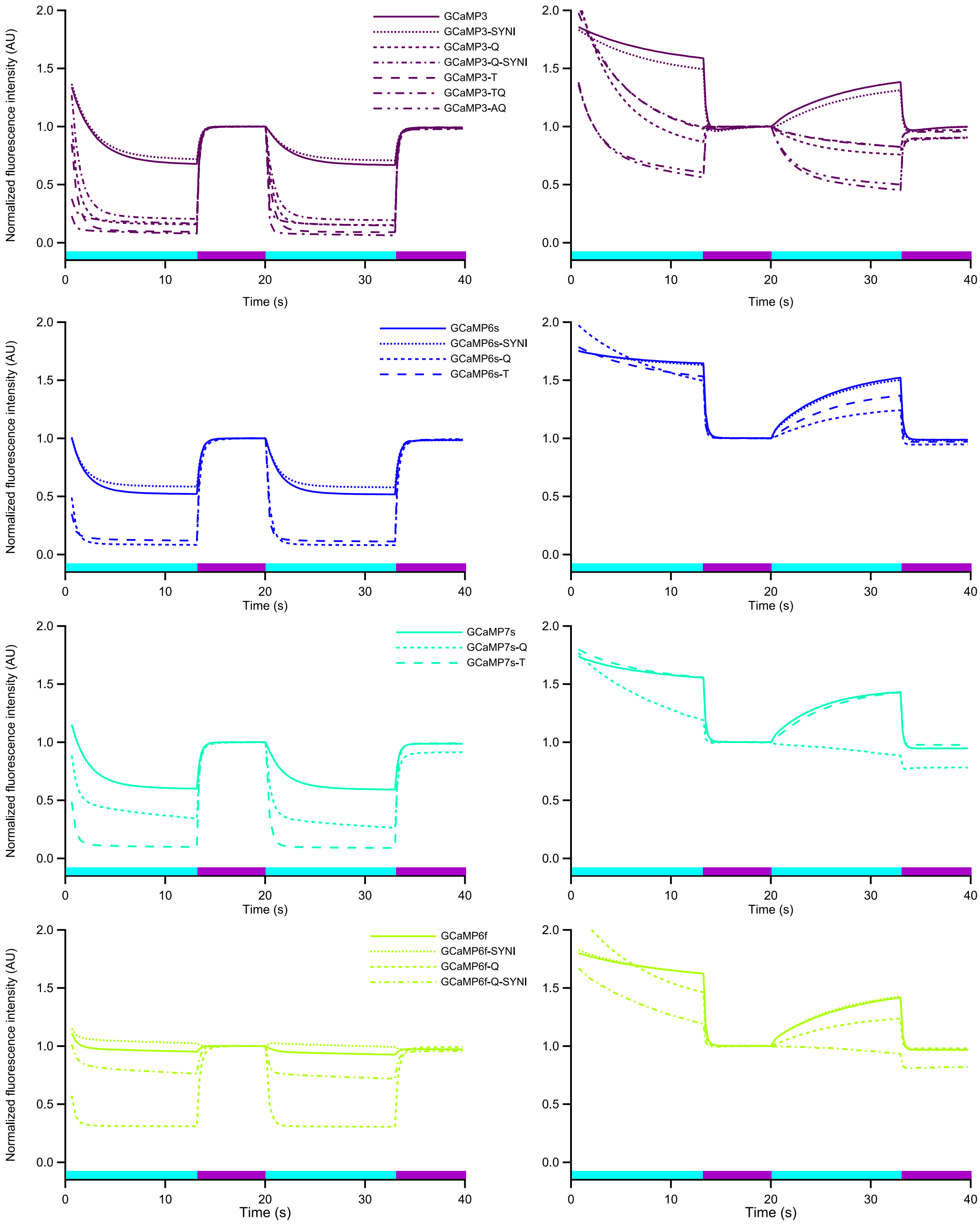


Supplementary Figure 2: Photoswitching experiments on rsGECIs in bacterial lysates supplemented with 5 mM $CaCl_2$ (left) and 1 mM EGTA (right). The cyan and violet highlights indicate the illumination sequence.

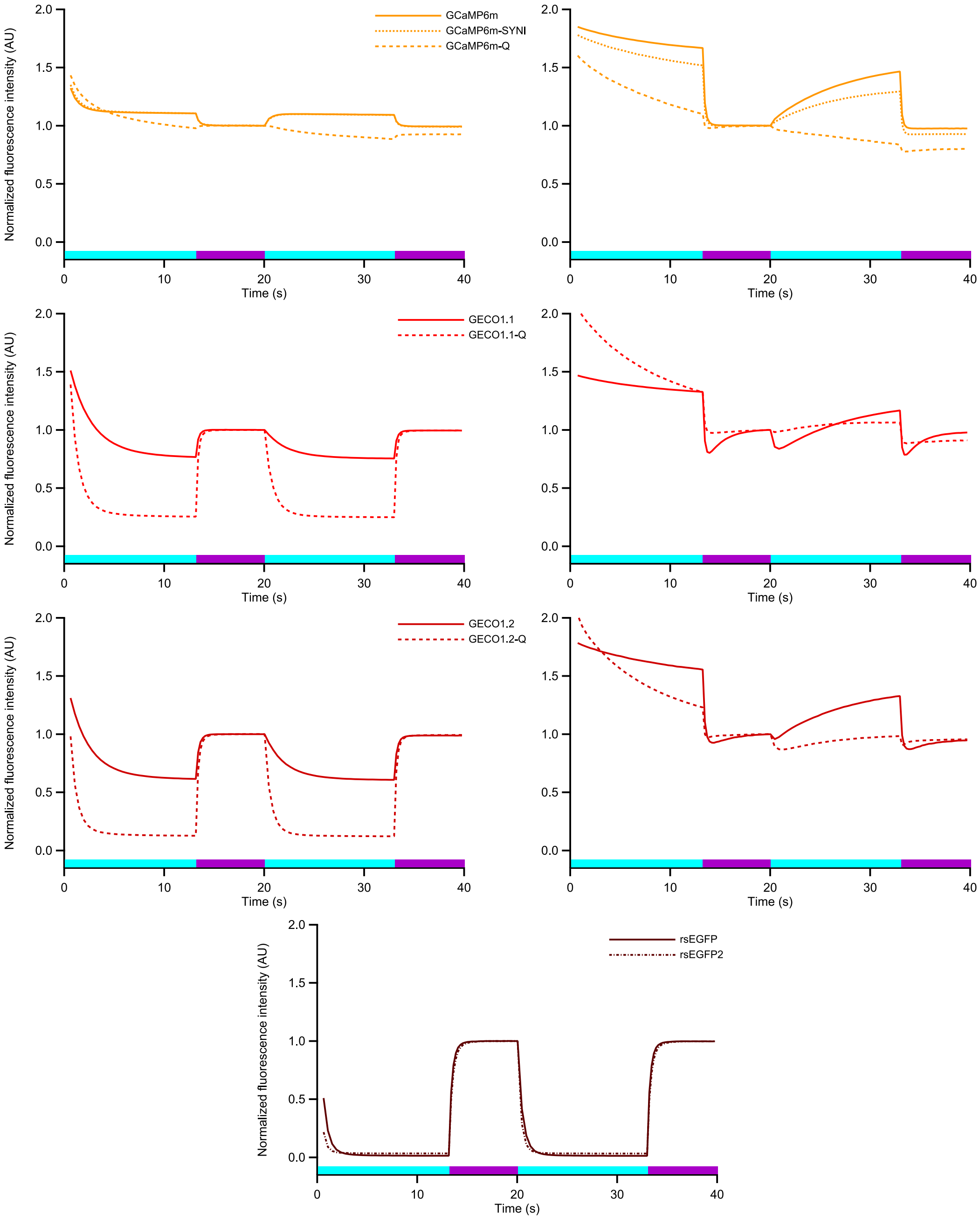


Supplementary Figure 3: Photoswitching experiments on rsGECIs in bacterial lysates supplemented with 5 mM $CaCl_2$ (left) and 1 mM EGTA (right). The cyan and violet highlights indicate the illumination sequence. rsEGFP and rsEGFP2 are shown as references for photochromic behavior.

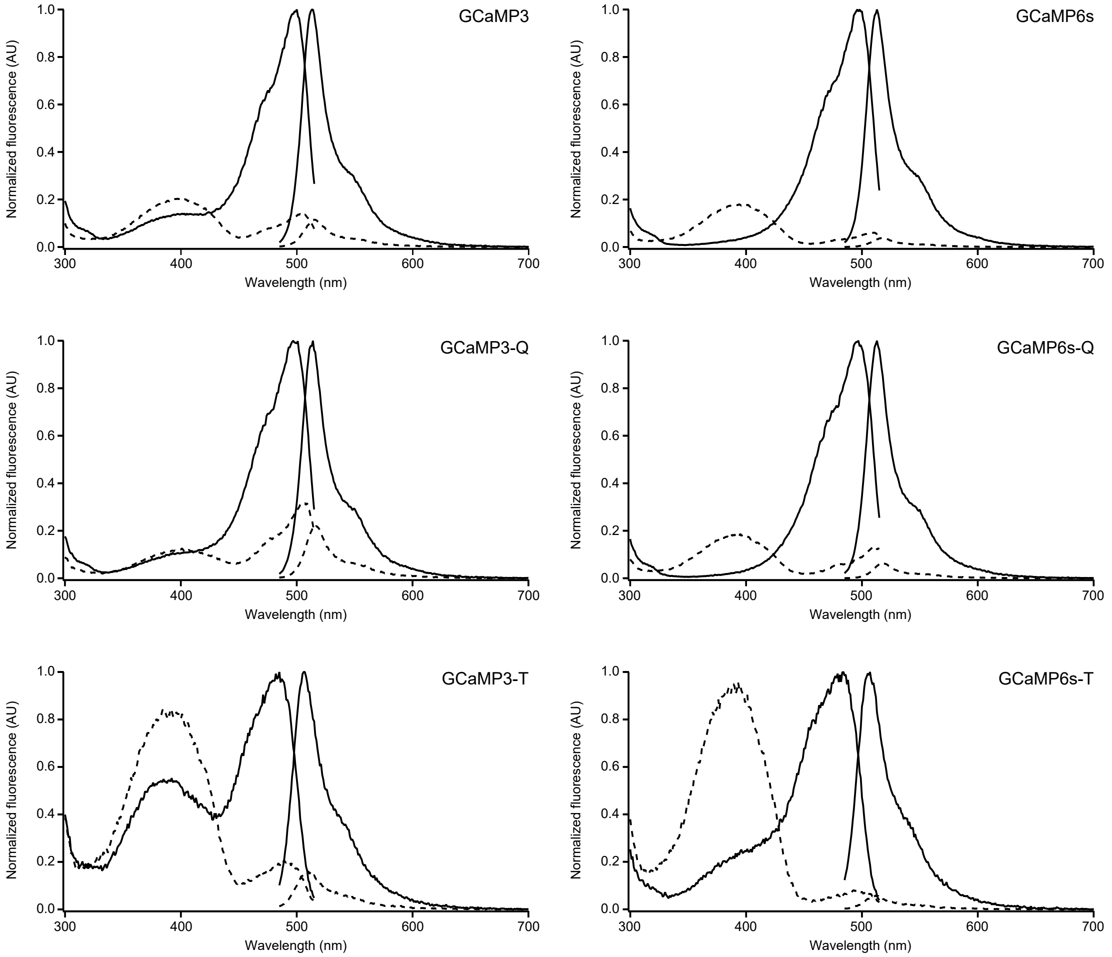


Supplementary Figure 4: Normalized excitation and emission spectra of the purified (rs)GECI proteins in their *apo* (dashed line) and *sat* (solid line) state.

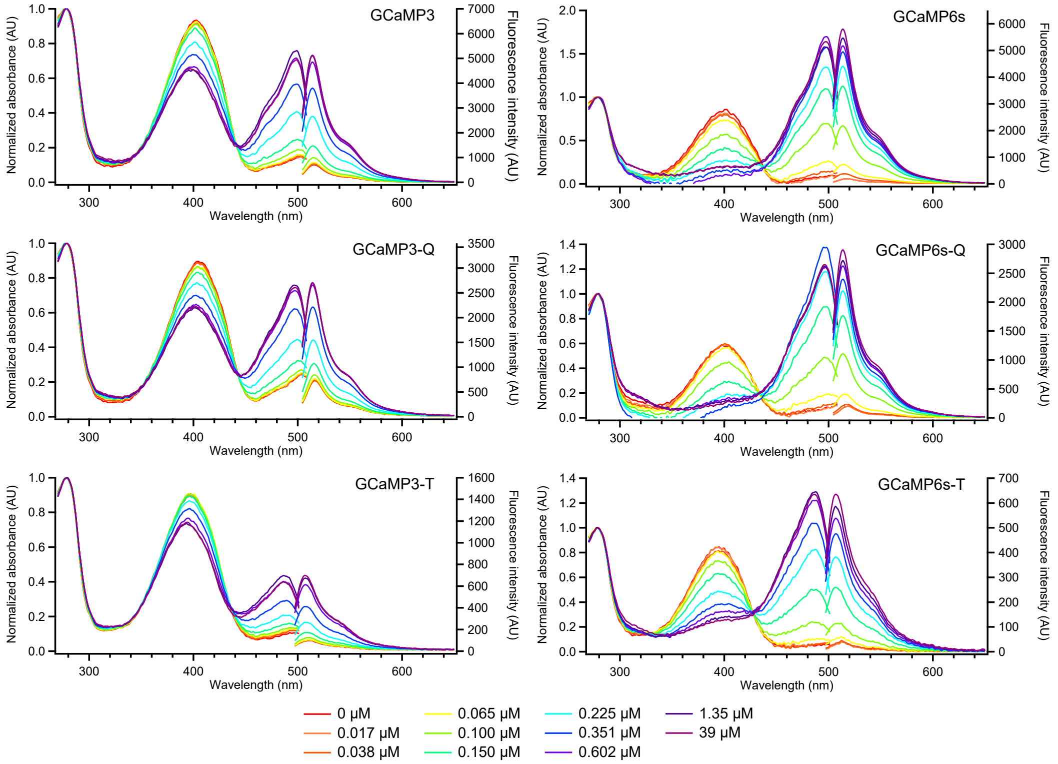


Supplementary Figure 5: Normalized absorbance and fluorescence emission spectra (exc. = 488 nm) of the purified (rs)GECI proteins in the fluorescent state for different concentrations of $Ca^{2+}$.

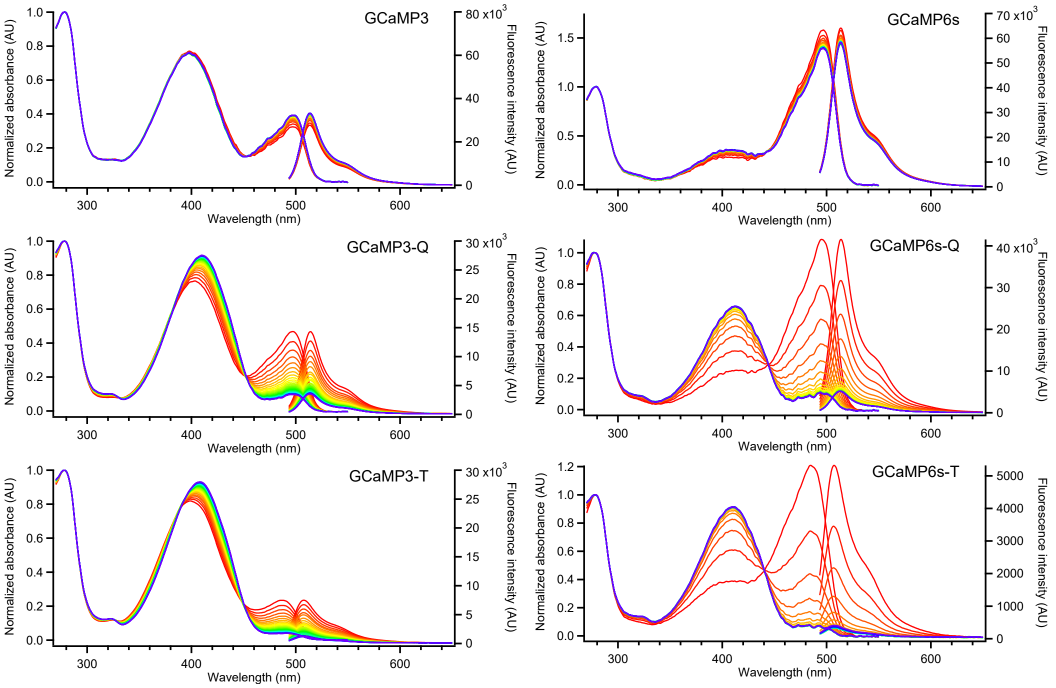


Supplementary Figure 6: Normalized absorbance and fluorescence emission spectra (exc. = 488 nm) of the purified (rs)GECI proteins at high $\left[Ca^{2+}\right]$ during cyan illumination. The proteins switch from the fluorescent on-state (red) to the non-fluorescent off-state (blue).

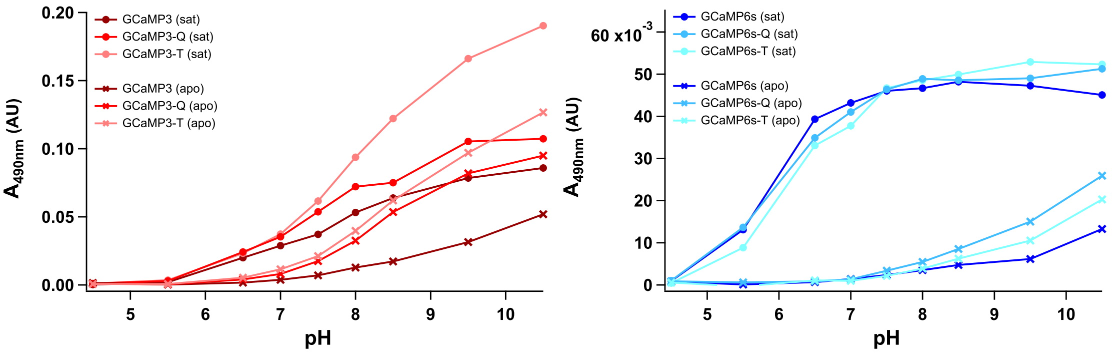


Supplementary Figure 7: pH dependence of the absorption maxima of the anionic form for the (rs)GECIs variants.

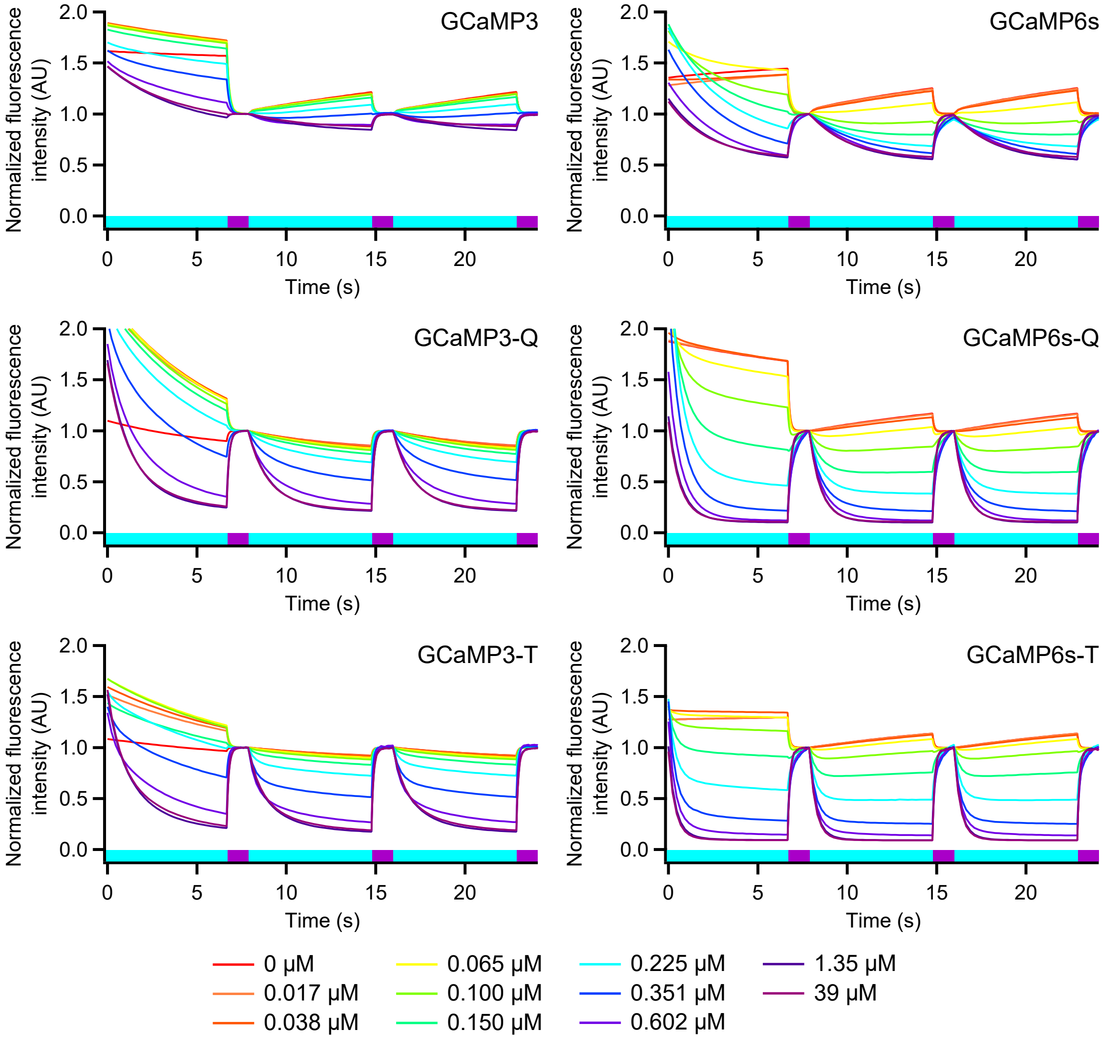


Supplementary Figure 8: $Ca^{2+}$ dependence of the photochromism of the 6 (rs)GECI variants.

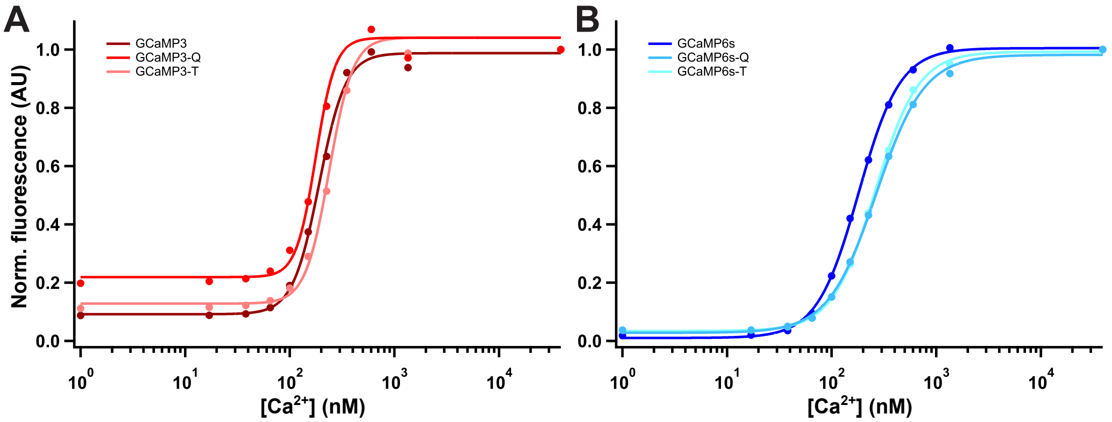


Supplementary Figure 9: $Ca^{2+}$ titration of the fluorescence of purified (rs)GECIs.

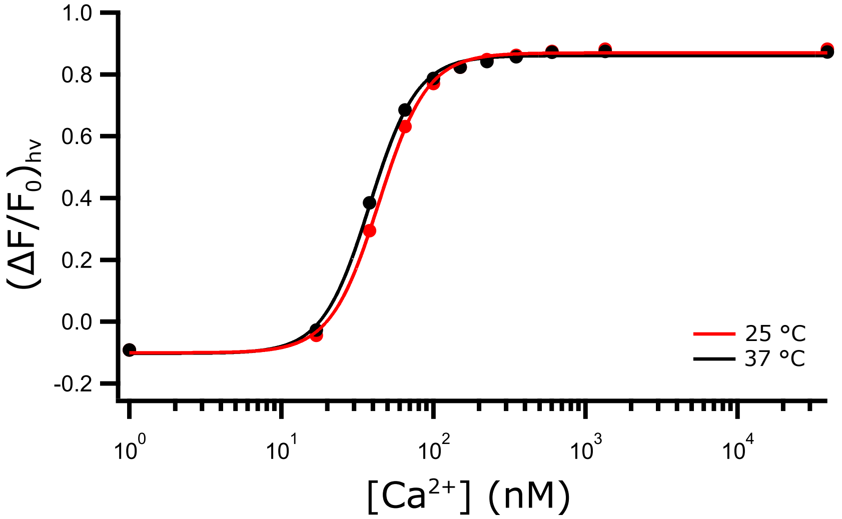


Supplementary Figure 10: $Ca^{2+}$ titrations of GCaMP6s-Q at room temperature (25°C) and 37°C.

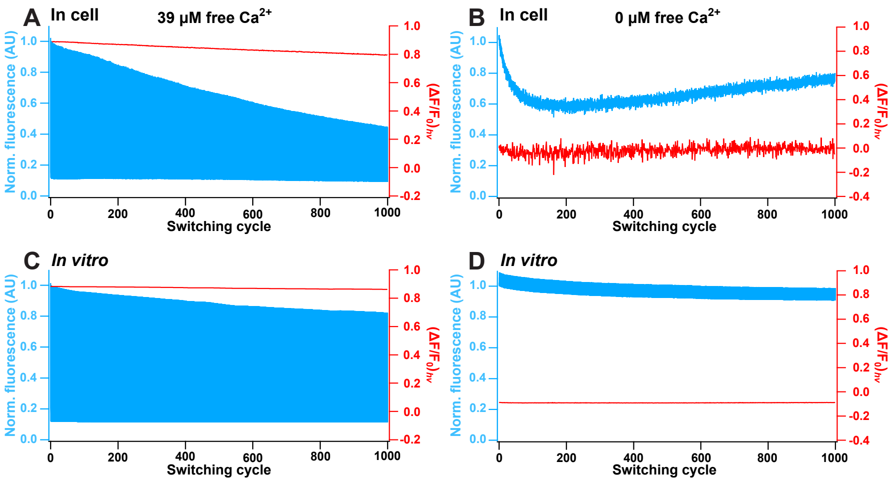


Supplementary Figure 11: Comparison between *in vitro* and in-cell fatigue measurements. (**A**) Fatigue measurement shown for one cell at high $Ca^{2+}$ concentration (*sat*). (**B**) Cell fatigue in calcium-free buffer (*apo*) shown for one cell. (**C**) *In vitro* fatigue at saturation $Ca^{2+}$ condition (*sat*). (**D**) *In vitro* fatigue in calcium-free buffer (*apo*). Cells were permeabilized as described in the materials and methods. Note that the high fatigue resistance *in vitro* also reflects the fact that the field-of-view of the imaging does not cover the full protein solution, meaning that replenishment with 'fresh' molecules occurs throughout the experiment.

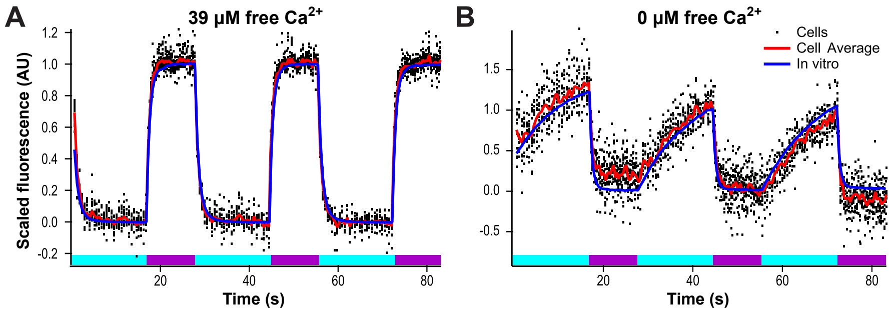


Supplementary Figure 12: Comparison of *in vitro* and in cell kinetics over three switching cycles under $Ca^{2+}$-saturing and $Ca^{2+}$-free conditions. The measured fluorescence values are offset and scaled to a range between 0 and 1. The black dots represent the measurements of switching cycles in different cells. The red trace show the average of the cells and the blue trace the *in vitro* trace. ($n = 22$ cells for *sat* and 14 for *apo*.)

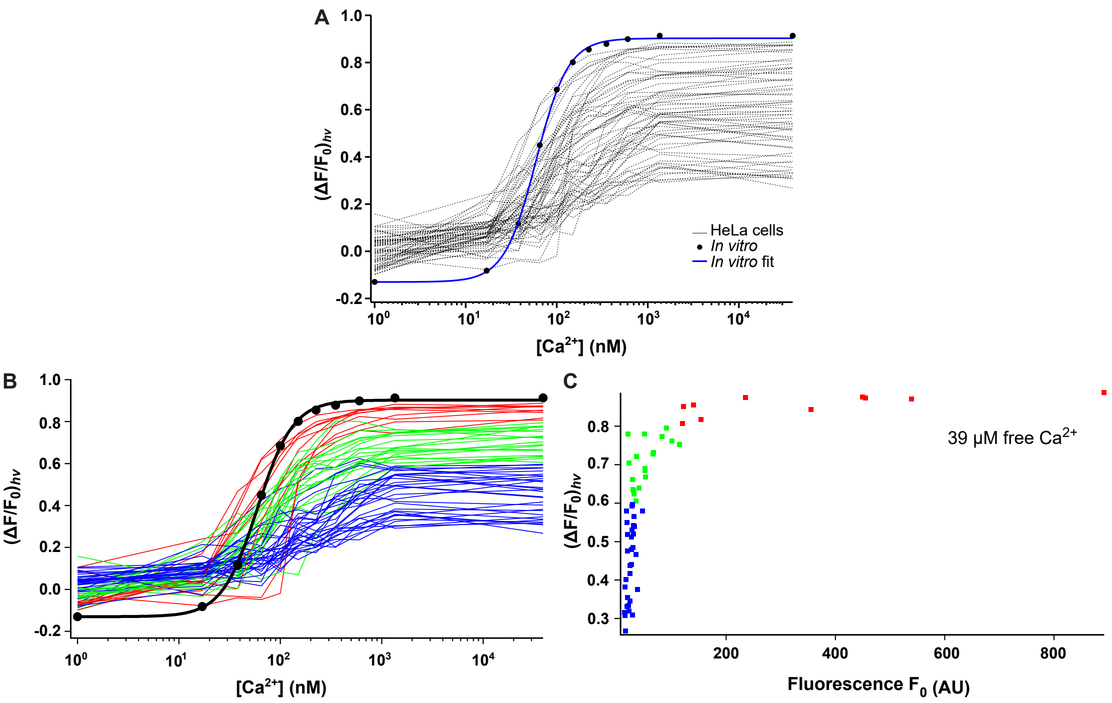


Supplementary Figure 13: (A) Titration curves of individual HeLa cells transfected with GCaMP6s-Q. All cells were washed with HHBSS(-), incubated in HHBSS(-) supplemented with ionomycin and EGTA for 10 minutes, and placed in Zero Free Calcium Buffer supplemented with ionomycin and saponin. The external $Ca^{2+}$ concentration was gradually increased following the reciprocal dilution method via medium exchange with 39 µM Free Calcium Buffer supplemented with ionomycin and saponin, as described in the method description of the *in vitro* $Ca^{2+}$ titrations. At every concentration, we assayed GCaMP6s-Q photoswitching and calculated $(\Delta F/F_0)_{h\nu}$ values. The HeLa titration curves are compared to the *in vitro* GCaMP6s-Q titration curve measured using the same settings (black markers and blue sigmoid fit). (B) Same data as shown in panel A but the traces have been color-coded according to cell brightness as shown in panel (C). ($n$ = 60 cells.)

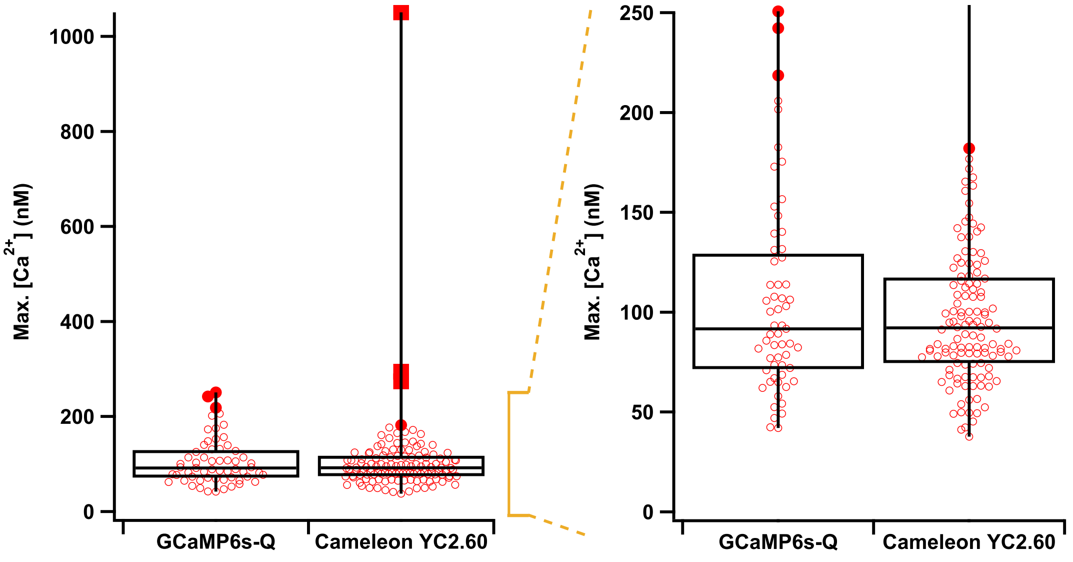


Supplementary Figure 14: Comparison of $Ca^{2+}$ levels in histamine-stimulated HeLa cells acquired using PEAQ with GCaMP-6sQ and the FRET based indicator Yellow Cameleon 2.60. The boxplots are indicating the median, the first and the third quartile. The whiskers are illustrating the maximum and minimum values. Filled red dots represent outliers (more than 1.5 times the interquartile range above or below the third and first quartile), and filled red squares are far outliers (more than three times the interquartile range above or below the third and first quartile). Expansion shows the $Ca^{2+}$ concentration range between 0 and 250 nM. ($n$ = 59 cells for PEAQ and 124 for FRET.)

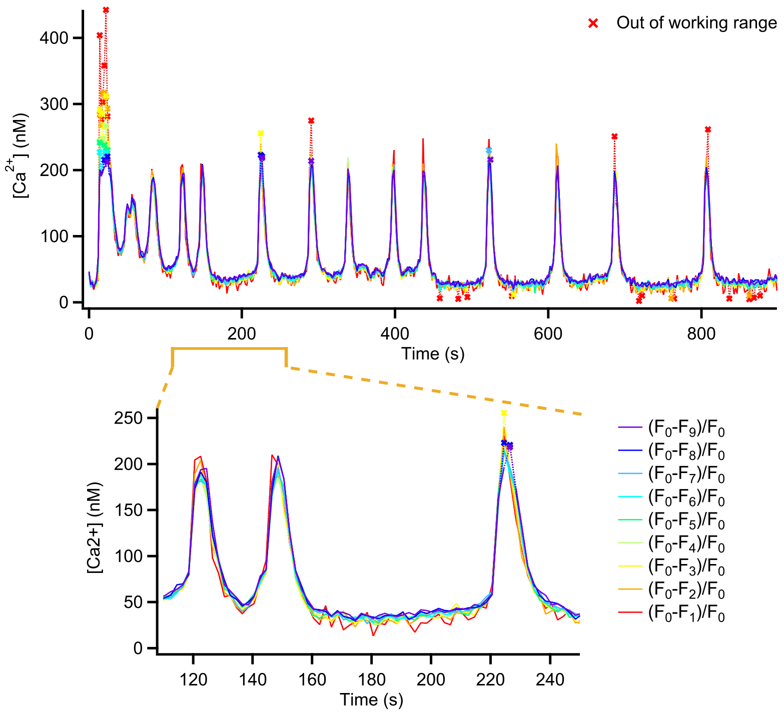


Supplementary Figure 15: $Ca^{2+}$ profiles calculated using different values for $F_{end}$ (see also Figure S16) for the data shown in main text Figure 3.

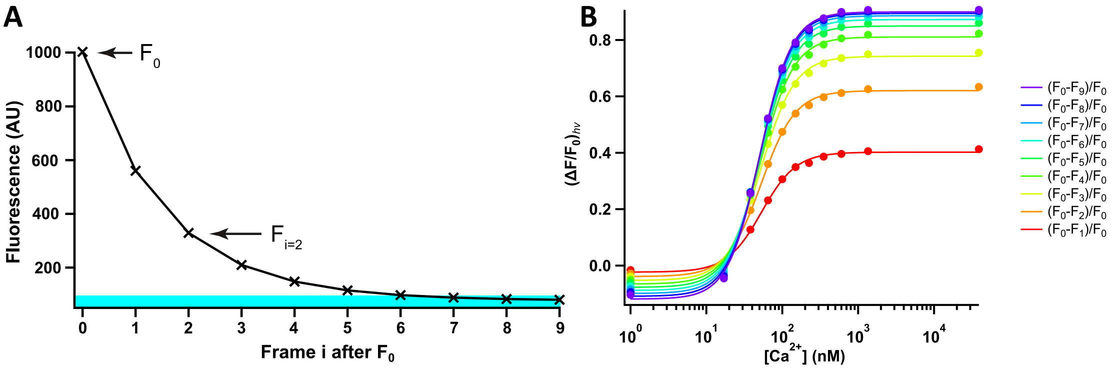


Supplementary Figure 16: (A) Illustration of the concept of using different $F_{end}$ images to calculate the $(\Delta F/F_0)_{h\nu}$ metric. Here the second image after $F_0$ is shown as an example. (B) *In vitro* titration curves for different definitions of $(\Delta F/F_0)_{h\nu} = (F_0 - F_i)/F_0$, where $F_i$ corresponds to the fluorescence intensity measured in image i after $F_0$.

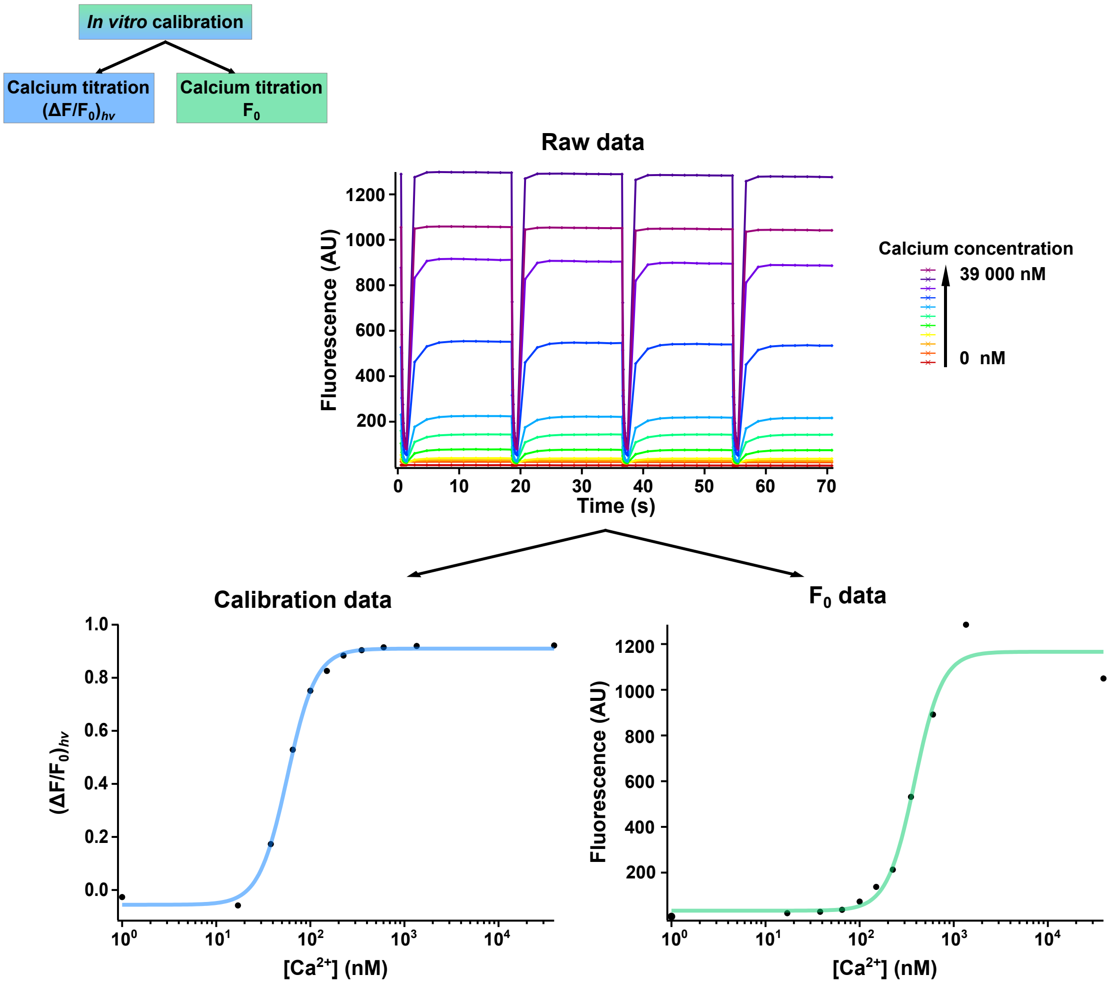


Supplementary Figure 17: *In vitro* titration of purified GCaMP6s-Q protein with an intermittent calibration imaging scheme that is identical to the scheme used to image histamine-induced $Ca^{2+}$-oscillations in transfected HeLa cells. Off/on switching calibration cycles are followed by eight $F_0$ measurements. The data is split into two subsets: one subset containing the switching cycles, that result in calibration $(\Delta F/F_0)_{h\nu}$ values correlated to the $[Ca^{2+}]$, and one set containing the $F_0$ data points. Both data sets are displayed in separate $Ca^{2+}$ titration curves, and are necessary to analyze the live cell data.

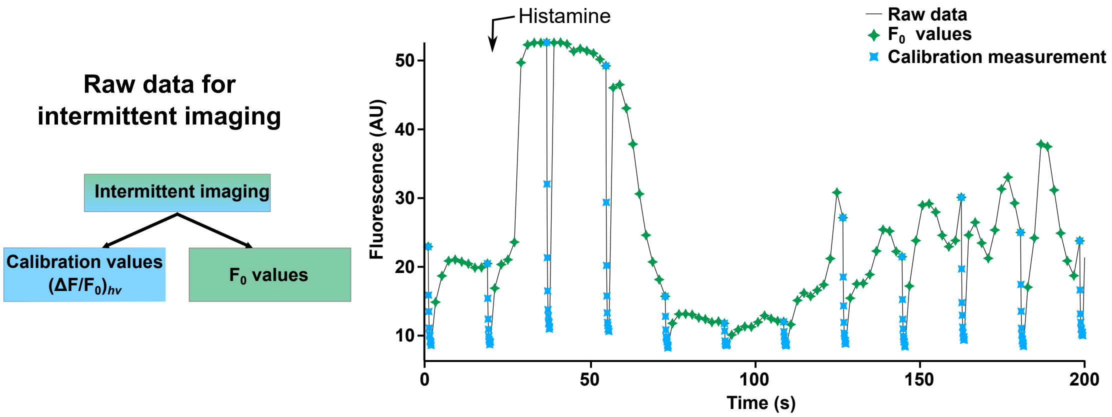


Supplementary Figure 18: Raw fluorescence data of a HeLa cell stimulated with histamine (25 μM), and imaged over 15 minutes using intermittent PEAQ (iPEAQ) biosensing. Only the initial response (200 s) is displayed. Each on/off switching calibration cycle (blue markers) is followed by eight $F_0$ measurements (green markers).

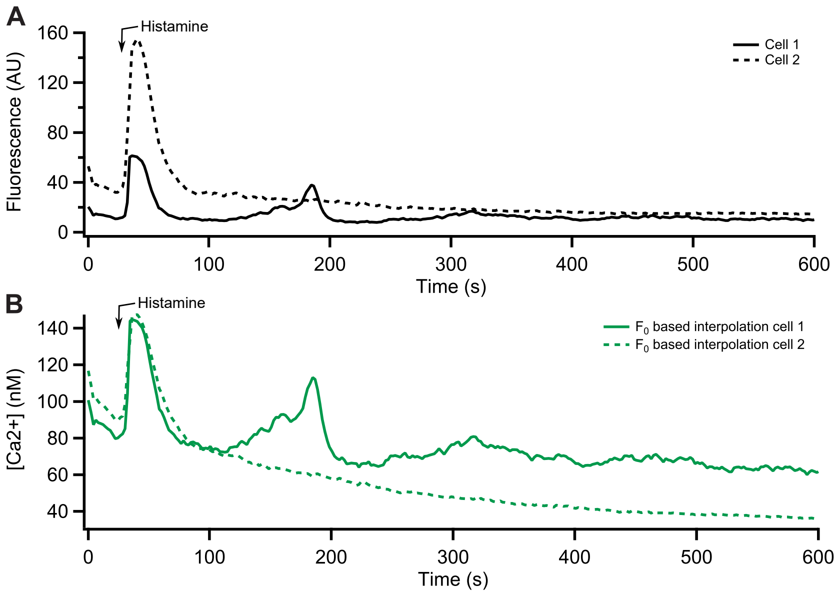


Supplementary Figure 19: Two HeLa cells stimulated with 25 μM histamine, imaged using intermittent PEAQ biosensing. (**A**) Raw fluorescence intensity traces based on the $F_0$ measurements. (**B**) $[Ca^{2+}]$ traces based on PEAQ biosensing.

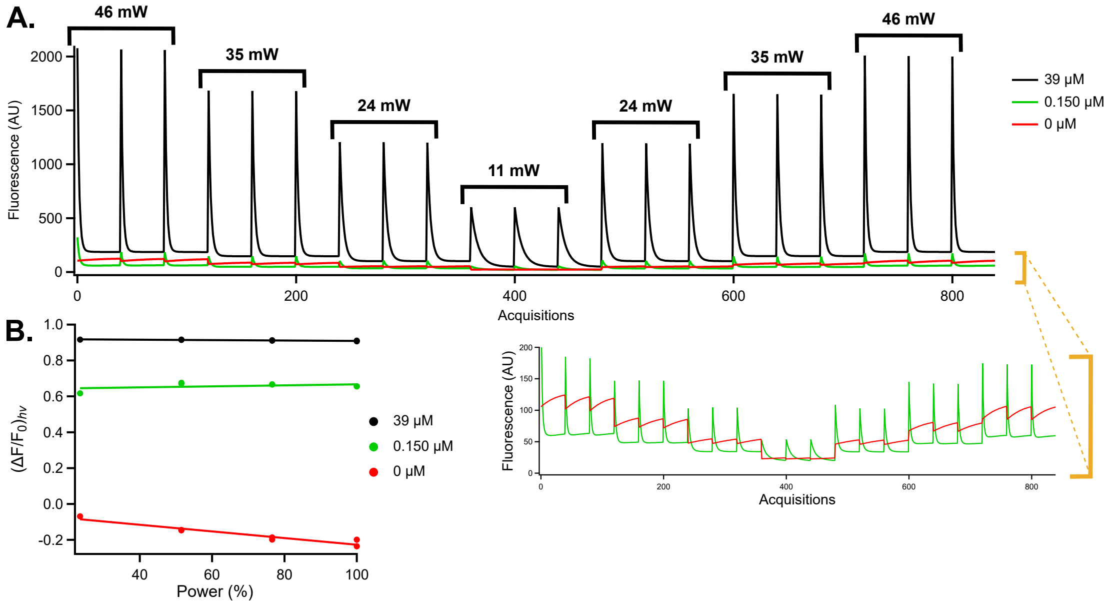


Supplementary Figure 20: *In vitro* verification of PEAQ biosensing power independence. (**A**) Raw intensity traces of purified GCaMP6s-Q protein, diluted in 0 mM, 5 mM, or 10 mM CaEGTA Buffer, resulting in 0 μM, 0.15 μM, or 39 μM free $Ca^{2+}$. At these three different concentrations, four different excitation powers were tested for off-switching. (**B**) $(\Delta F/F_0)_{h\nu}$ values calculated for the four different excitation powers.

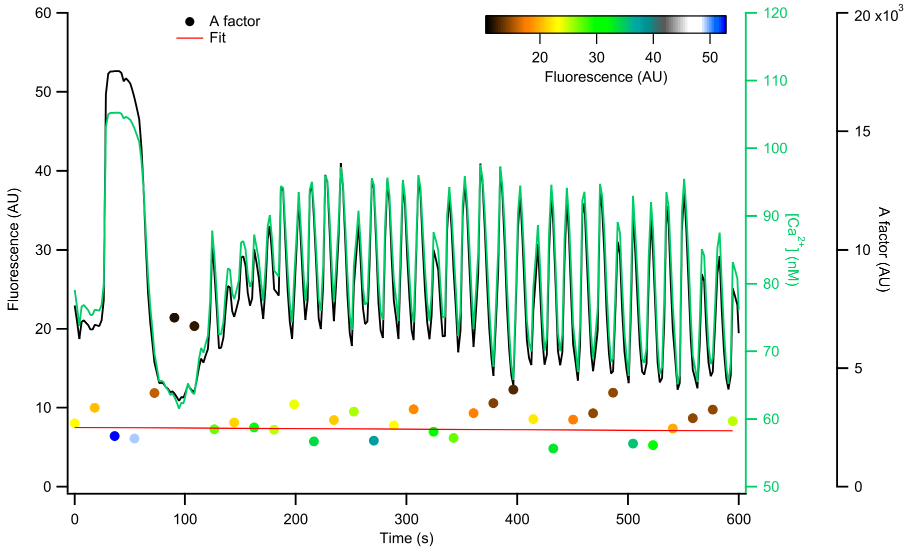


Supplementary Figure 21: iPEAQ data showing the values of the raw *A* correction factors (explained in the materials and methods) for the measurement shown in main text Figure 4D, as well as the fit used to calculate interpolated values for this factor.

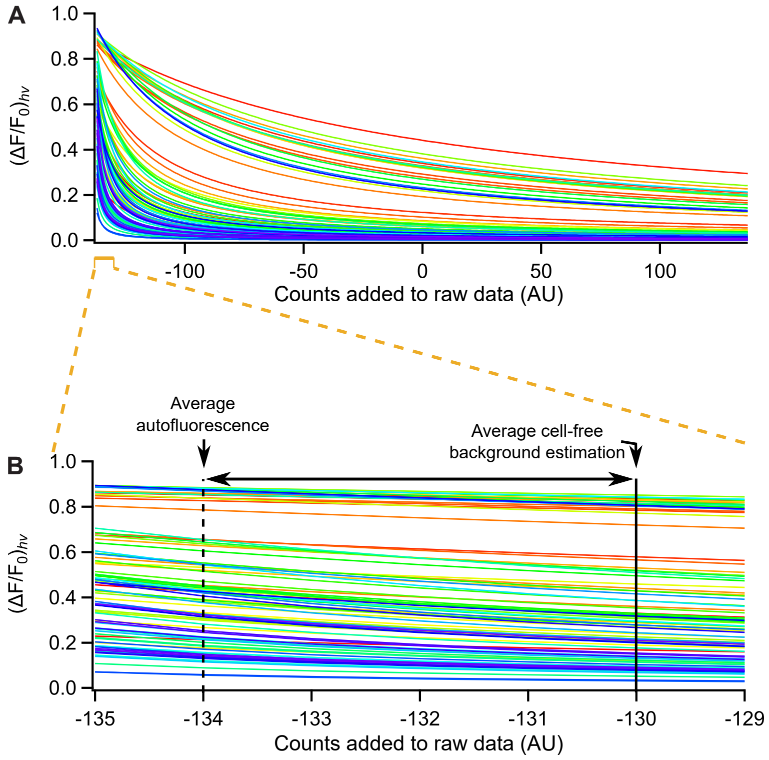


Supplementary Figure 22: (A) Calculated $(\Delta F/F_0)_{h\nu}$ values for varying amounts of counts added to the raw measured cell fluorescence. A value of zero means that the raw data is used as measured, that is, without correction of any kind. The different traces show the behaviour of different $(\Delta F/F_0)_{h\nu}$ measurements for the data shown in Figure 3D and E. (B) Expansion showing the region for a subtraction of 130 counts, which is a typical background estimate based on the measured signal of a cell-free region. The gray region shows the intensity of the estimated autofluorescence based on the measurement of untransfected cells.
